# Catalytic rewiring of RuvC-II catalytic site activates *trans*-cleavage in Fanzor2

**DOI:** 10.64898/2026.06.23.733943

**Authors:** Bingrong Xu, Xiaoqiang Huang, Xiaosong Han, Shengsong Xie, Xinyun Li, Jianlin Han, Di Wu, Shuaicheng Li, Shuhong Zhao, Changzhi Zhao

## Abstract

Fanzors are eukaryotic RNA-guided endonucleases that mediate programmable *cis*-cleavage in eukaryotic cells, but their potential target-induced collateral (*trans*) cleavage activity remains largely unexplored. Here, we show that catalytic rewiring of the RuvC-II catalytic site activates robust *trans*-cleavage activity in *Acanthamoeba polyphaga* mimivirus Fanzor2 (ApmFz2). The representative variant ApmFz2-EP displayed DNA– and RNA-triggered *trans*-cleavage, attenuated *cis*-cleavage, minimal TAM dependence, and activation by as few as seven nucleotides of guide-target complementarity. Mechanistic analyses indicate that relieving steric constraints surrounding the conserved alternative glutamate within the RuvC-II activates *trans*-cleavage and reshapes target-recognition specificity. Coupling ApmFz2-EP with nucleic acid amplification enabled FINDER, a Fanzor-based diagnostic platform for sensitive pathogen detection and broad detection across genetically diverse target subtypes. In addition, the mismatch-sensitive ApmFz2-EA variant enabled specific single-nucleotide variant (SNV) genotyping. Together, this work expands the functional scope of Fanzors and identifies catalytic-center engineering as a strategy for developing compact, programmable nucleic acid diagnostics.

## Introduction

Programmable nucleic acid detection platforms based on RNA-guided nucleases have transformed molecular diagnostics^1–4^. CRISPR effectors such as Cas12a and Cas13a mediate target-triggered collateral (*trans*) cleavage of single-stranded DNA (ssDNA) or single-stranded RNA (ssRNA) reporters, enabling sensitive detection of pathogens and genetic variants^5–7^. Despite their broad use, many Cas12a systems require T-rich protospacer adjacent motifs (PAMs), which can restrict target accessibility and limit diagnostic flexibility^8,9^. Recent evolutionary studies suggest that obligate mobile element-guided activity (OMEGA) systems are thought to represent the evolutionary ancestors of Cas9 and Cas12 effectors^10,11^. Among these, TnpB proteins encoded by IS200/IS605 transposons are widely regarded as evolutionary precursors of Cas12 effectors^12,13^. Several TnpB homologs also exhibit target-triggered *trans*-cleavage activity^14,15^, indicating that collateral cleavage can occur across distinct classes of RNA-guided nucleases.

Fanzors are recently discovered eukaryotic RNA-guided nucleases encoded by mobile genetic elements and distributed across giant viruses and diverse eukaryotic lineages, including metazoans, fungi, and protists^16–18^. Fanzors use ωRNAs to direct programmable DNA cleavage and retain a conserved RuvC nuclease core that shares high structural similarity with those of their prokaryotic homologs, TnpB and Cas12^16,19^. Recent studies have demonstrated their genome-editing activity in eukaryotic cells^16,20–22^. TnpB is also thought to be the evolutionary ancestor of Fanzor proteins. Despite their evolutionary relationship with TnpB, naturally occurring Fanzors generally show weak or undetectable *trans*-cleavage activity^18,19^.

A key structural distinction between these TnpB and Fanzor nucleases lies within the catalytic center. In canonical TnpBs proteins, the RuvC-II motif contains a conserved catalytic glutamate. In Fanzors, this residue is displaced toward the C terminus, generating a conserved alternative glutamate^17,18,23^. This unusual active-site architecture raises the possibility that evolutionary remodeling of the catalytic center contributed to the loss or attenuation of *trans*-cleavage activity in Fanzors.

Here, we show that reconstructing the canonical “D-E-D” catalytic configuration in ApmFz2 activates robust *trans*-cleavage activity while attenuating canonical *cis*-cleavage. Systematic mutational analyses further reveal that side-chain volume at the highly conserved alternative glutamate position regulates both *trans*-cleavage activation and target-recognition specificity, generating variants that range from broad mismatch tolerance to single-nucleotide discrimination. By using these properties, we developed ApmFz2-based diagnostic platforms for sensitive pathogen detection and accurate single-nucleotide variant (SNV) genotyping. Together, our findings identify catalytic-site evolution as a determinant of ApmFz2 function and establish engineered Fanzors as programmable diagnostic effectors.

## Results

### RuvC-II active-site rewiring activates ApmFz2 *trans*-cleavage

TnpB proteins encoded by IS200/IS605 transposons have been shown to possess target-dependent *trans*-cleavage activity^14,15^. As the eukaryotic homologs of TnpB, we reasoned that Fanzors might retain latent collateral cleavage potential. Unlike canonical TnpB proteins, which contain a conserved “D-E-D” catalytic triad, Fanzor possesses a noncanonical RuvC-II motif in which the catalytic glutamate is replaced by an inert proline or glycine^18,23^ (Fig. 1a). Instead, these systems harbor a conserved alternative glutamate downstream of the canonical RuvC-II position, which is thought to compensate functionally for the missing RuvC-II glutamate and establish a noncanonical “D-altE-D” catalytic configuration^17,18^. We therefore hypothesized that relocating the alternative conserved glutamate to the canonical RuvC-II glutamate position, thereby restoring the canonical “D-E-D” configuration, could reconfigure the catalytic center and promote *trans*-cleavage activity in Fanzor nucleases.

**Fig. 1.**
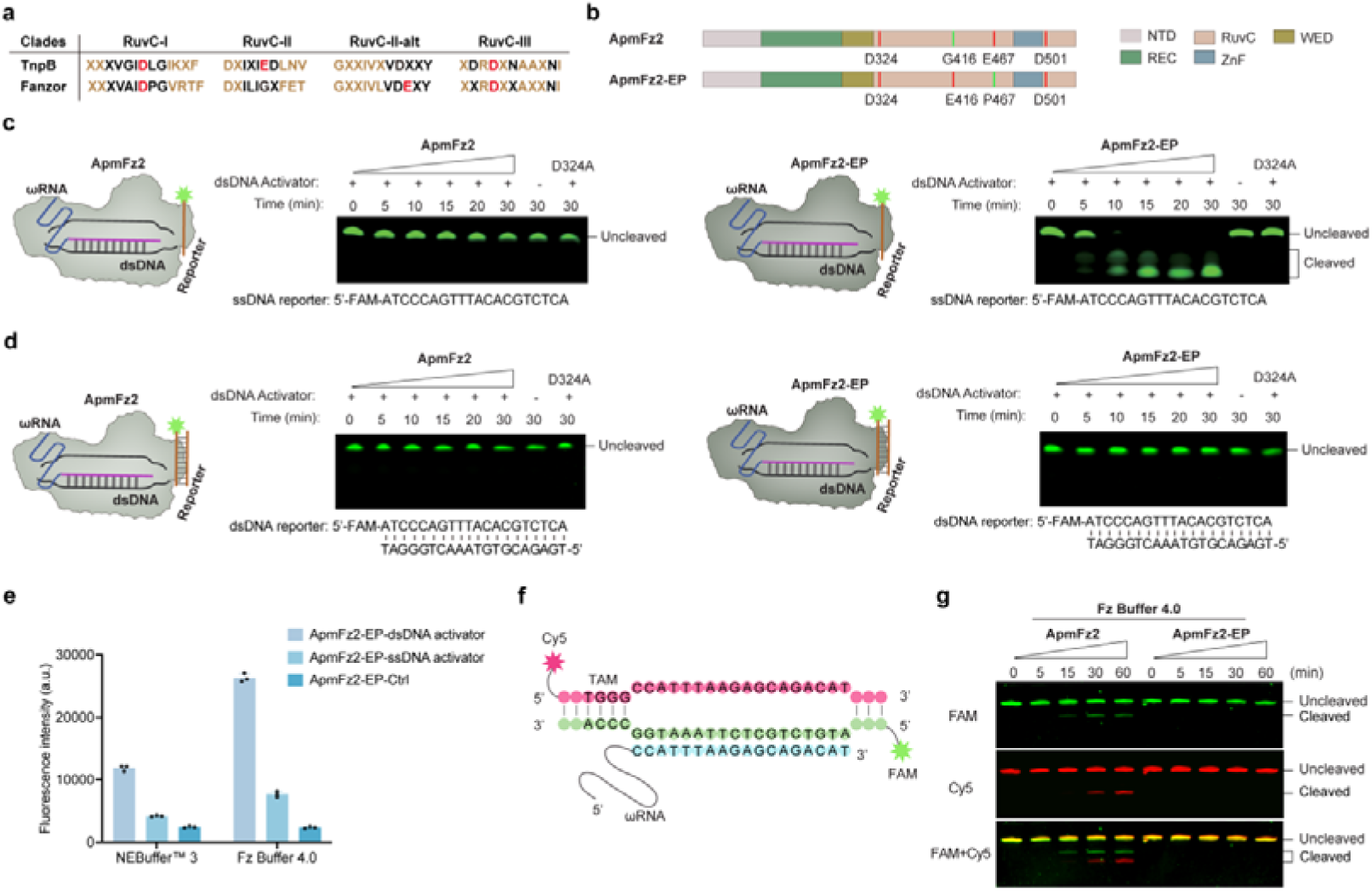
Evaluation of ApmFz2-EP *trans*– and *cis*-cleavage activity. **a**, Alignment of TnpB and Fanzor clade consensus sequences across the RuvC-I, RuvC-II, and RuvC-III motifs. **b**, Domain architectures of wild-type ApmFz2 and engineered ApmFz2-EP. ApmFz2-EP was generated by introducing G416E and E467P substitutions into ApmFz2. **c,d**, *Trans*-cleavage activity of ApmFz2 and ApmFz2-EP toward 5′-FAM-labeled ssDNA (c) or dsDNA (d) reporters in the presence of a dsDNA activator, analyzed by denaturing gel electrophoresis. **e**, Comparison of ApmFz2-EP *trans*-cleavage activity in NEBuffer™ 3 and Fz Buffer 4.0 after activation by dsDNA or ssDNA activators. Ctrl, no-template control. Data are shown as mean ± s.d. (*n* = 3 technical replicates). **f**, Schematic of the dsDNA substrate labeled with Cy5 and FAM at opposite termini for *cis*-cleavage analysis. **g,** *Cis*-cleavage activity of ApmFz2 and ApmFz2-EP on the dsDNA substrate shown in (f), analyzed by denaturing gel electrophoresis. ApmFz2, ApmFz2-EP, ApmFz2_D324A, and ApmFz2-EP_D324A proteins were purified using the pCold-TF expression system.

To test this hypothesis, we engineered a Fanzor2 homolog from *Acanthamoeba polyphaga* mimivirus (ApmFz2) by restoring the canonical RuvC-II glutamate (G416E) and replacing the alternative conserved glutamate with proline (E467P), the most frequent residue at the corresponding position in TnpB proteins^18,23^. This double mutant was designated ApmFz2-EP (Fig. 1b). ApmFz2-EP was expressed in *Escherichia coli* using the pCold-TF expression system and purified to high purity, as confirmed by SDS–PAGE analysis (Extended Data Fig. 1). We first assessed whether ApmFz2-EP could mediate *trans*-cleavage of an ssDNA reporter after activation by double-stranded DNA (dsDNA) activators. ApmFz2-EP showed robust *trans*-cleavage activity against the ssDNA reporter, whereas wild-type AmpFz2 showed minimal activity, comparable to the catalytically inactive RuvC-I mutants ApmFz2_D324A (Fig. 1c). We next evaluated *trans*-cleavage activity against dsDNA reporters in the presence of dsDNA activators. In contrast to ssDNA reporters, dsDNA reporters were barely cleaved by either ApmFz2-EP or wild-type ApmFz2 (Fig. 1d), indicating that ApmFz2-EP preferentially mediates *trans*-cleavage of ssDNA rather than dsDNA reporters, consistent with the substrate preferences reported for Cas12a, Cas12b, and TnpB nucleases^4,6,15^.

We next optimized the reaction conditions to improve ApmFz2-EP *trans*-cleavage activity. NEBuffer™ 3 was initially tested as a reference condition but showed suboptimal performance. We therefore developed a custom formulation, termed Fz Buffer 1.0, which enhanced *trans*-cleavage activity relative to NEBuffer™ 3 (Extended Data Fig. 2a,f). We then systematically evaluated pH, divalent metal ion species and concentrations, DTT concentration, and KCl concentration, ultimately identifying Fz Buffer 4.0 as an optimized formulation that markedly enhanced ApmFz2-EP *trans*-cleavage activity (Fig. 1e and Extended Data Fig. 2b-f). Previous studies have shown that *trans*-cleavage activities in Cas12 and TnpB systems are preferentially activated by ssDNA activators^15,24^. We therefore compared the ability of ssDNA and dsDNA activators to stimulate ApmFz2-EP *trans*-cleavage activity. Unexpectedly, ApmFz2-EP was activated more efficiently by dsDNA activators than by ssDNA activators (Fig. 1e and Extended Data Fig. 3).

Cas12 and TnpB nucleases use a single RuvC domain to mediate RNA-guided, site-specific *cis*-cleavage of target dsDNA and *trans*-cleavage of nonspecific ssDNA substrates^12,13,25^. Fanzors also possess a RuvC nuclease domain that mediates RNA-guided dsDNA cleavage, and the conserved alternative glutamate has been reported to be required for efficient nuclease activity^18^. These observations raised the possibility that rewiring the noncanonical Fanzor RuvC active site may promote *trans*-cleavage activity at the expense of canonical *cis*-cleavage. To evaluate ApmFz2-EP *cis*-cleavage activity, we used a dsDNA target labeled with Cy5 and FAM at opposite termini (Fig. 1f). ApmFz2-EP showed minimal cleavage of the dsDNA substrate, whereas wild-type ApmFz2 retained robust *cis*-cleavage activity (Fig. 1g). Together, these findings indicate that rewiring the noncanonical Fanzor catalytic center functionally reprograms ApmFz2 into a *trans*-cleavage-active nuclease and conferring preferential activation by dsDNA activators, while substantially impairing canonical *cis*-cleavage.

### Characterization of ApmFz2-EP *trans*-cleavage

ApmFz2 has been reported to exhibit optimal *cis*-cleavage activity at 30-40 °C *in vitro*^18,19^. To define the temperature range for ApmFz2-EP *trans*-cleavage, we monitored reporter cleavage across temperatures from 25 to 55 °C using a fluorophore-quencher-labeled ssDNA reporter (Fig. 2a). ApmFz2-EP showed maximal *trans*-cleavage activity at 35 to 45 °C, whereas fluorescence signals were substantially reduced at lower (25 °C) or higher (50 °C) temperatures. These results suggest that catalytic rewiring does not substantially alter the temperature dependence of ApmFz2-EP activity *in vitro*.

**Fig. 2.**
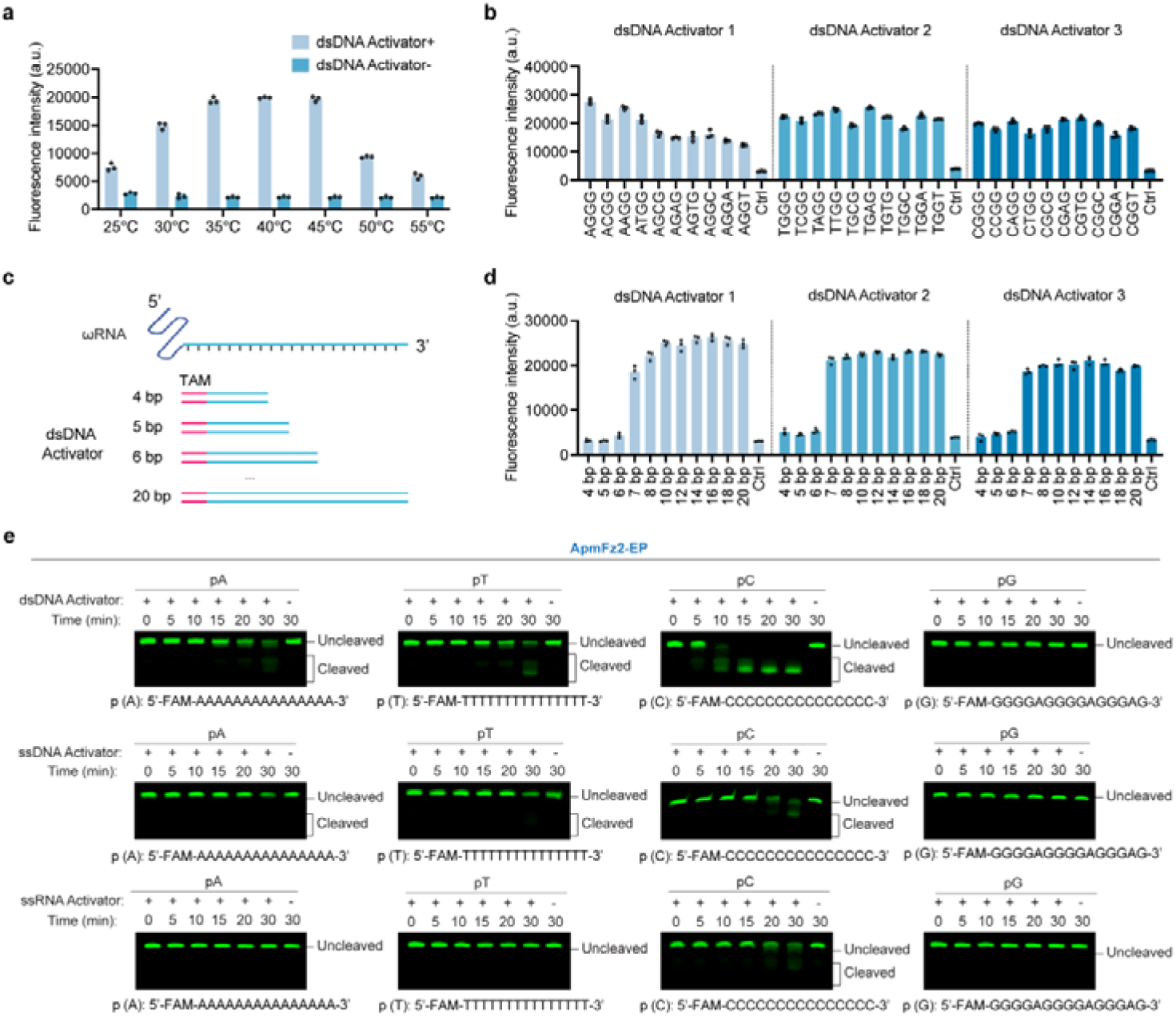
Characterization of ApmFz2-EP *trans*-cleavage. **a**, Temperature dependence of ApmFz2-EP *trans*-cleavage activity measured using a fluorescent ssDNA reporter. Data are shown as mean ± s.d. (*n* = 3 technical replicates). **b**, TAM requirements for ApmFz2-EP *trans*-cleavage activation, assessed using three dsDNA activators carrying 30 mutated TAM sequences. Ctrl, no-template control. Data are shown as mean ± s.d. (*n* = 3 technical replicates). **c**, Schematic showing stepwise truncation of a 20-bp dsDNA activator to determine the minimal guide-target pairing length required for *trans*-cleavage activation. **d**, Determination of the minimal guide-target pairing length required for ApmFz2-EP *trans*-cleavage activation using three dsDNA activators. Ctrl, no-template control. Data are shown as mean ± s.d. (*n* = 3 technical replicates). **e**, Reporter substrate preference of ApmFz2-EP *trans*-cleavage activity assessed using 5′-FAM-labeled poly(A), poly(T), poly(C), and poly(G) ssDNA reporters in the presence of dsDNA, ssDNA, or ssRNA activators, analyzed by denaturing gel electrophoresis. p(A), poly(A); p(T), poly(T); p(C), poly(C); p(G), poly(G). ApmFz2-EP protein was purified using the pCold-TF expression system.

Previous target-adjacent motif (TAM) depletion analysis showed that NGGG TAMs are required for ApmFz2 *cis*-cleavage^18^. To examine the TAM requirement for ApmFz2-EP *trans*-cleavage, we performed *trans*-cleavage assays using three dsDNA activators containing 30 mutated TAM sequences (Fig. 2b). All mutated TAM sequences efficiently activated *trans*-cleavage activity, indicating that ApmFz2-EP *trans*-cleavage does not strictly depend on specific TAM sequences.

We next defined the minimal guide-target pairing length required for ApmFz2-EP *trans*-cleavage activation. To do this, we designed a series of dsDNA activators containing complementary regions ranging from 4 to 20 bp (Fig. 2c). As few as seven nucleotides of guide-target complementarity were sufficient to activate ApmFz2-EP *trans*-cleavage activity, whereas target DNAs with fewer than seven complementary nucleotides failed to trigger detectable activity (Fig. 2d). This requirement differs substantially from those reported for Cas12a (≥14 nt) and TnpB nucleases (≥12 nt)^15,26^. We then profiled the substrate preference of ApmFz2-EP using homopolymeric FAM-labeled ssDNA reporters in the presence of ssDNA, dsDNA, or ssRNA activators. All three activator types induced ApmFz2-EP *trans*-cleavage activity (Fig. 2e). Both ssDNA and dsDNA activators induced broad reporter cleavage, with a strong preference for poly(C) reporters, weaker activity toward poly(A) and poly(T), and minimal cleavage of poly(G) substrates. Consistent with our earlier observations, ssDNA-mediated activation produced substantially weaker *trans*-cleavage activity than dsDNA-mediated activation (Fig. 1e and Extended Data Fig. 3). In contrast, ssRNA activators supported detectable cleavage only for poly(C) reporters, resembling the RNA-triggered reporter preference previously observed in Cas12f1 systems^27^.

Finally, we examined whether secondary structure influences ApmFz2-EP *trans*-cleavage activity using structured ssDNA reporters (Extended Data Fig. 4a). The activity profile closely paralleled that observed with linear reporters, with Stem-10C generating the strongest fluorescence signal (Extended Data Fig. 4b). These results further support a preference for C-rich ssDNA reporters. Together, these findings establish ApmFz2-EP as a target-dependent *trans*-cleavage nuclease that responds to both DNA and RNA inputs while preferentially cleaving C-rich ssDNA reporters.

### Small-to-medium side-chain substitutions at position 467 activate ApmFz2 *trans*-cleavage

The robust *trans*-cleavage activity of ApmFz2-EP following restoration of the RuvC-II glutamate (G416E) and substitution of the native Glu467 with proline (E467P) suggested that a less bulky amino acid at position 467 may be critical for *trans*-cleavage activation. To systematically assess the contribution of residue identity at this position, we replaced Pro467 in ApmFz2-EP with each of the other 18 amino acids, excluding the original glutamate, to generate a panel of ApmFz2-EX double variants (Fig. 3a). *Trans*-cleavage assays showed that variants carrying small-to-medium side-chain substitutions at position 467, including ApmFz2-EP, EV, EG, EC, EN, ET, ES, ED, and EA, exhibited robust *trans*-cleavage activity (Fig. 3b). In contrast, variants containing bulky substitutions displayed little or no detectable activity, except for ApmFz2-EH, which retained weak activity (Fig. 3b). These findings indicate that smaller side chains at position 467 favor *trans*-cleavage activation, potentially by creating additional space within the active-site region and allowing proper positioning of the reconstructed G416E residue.

**Fig. 3.**
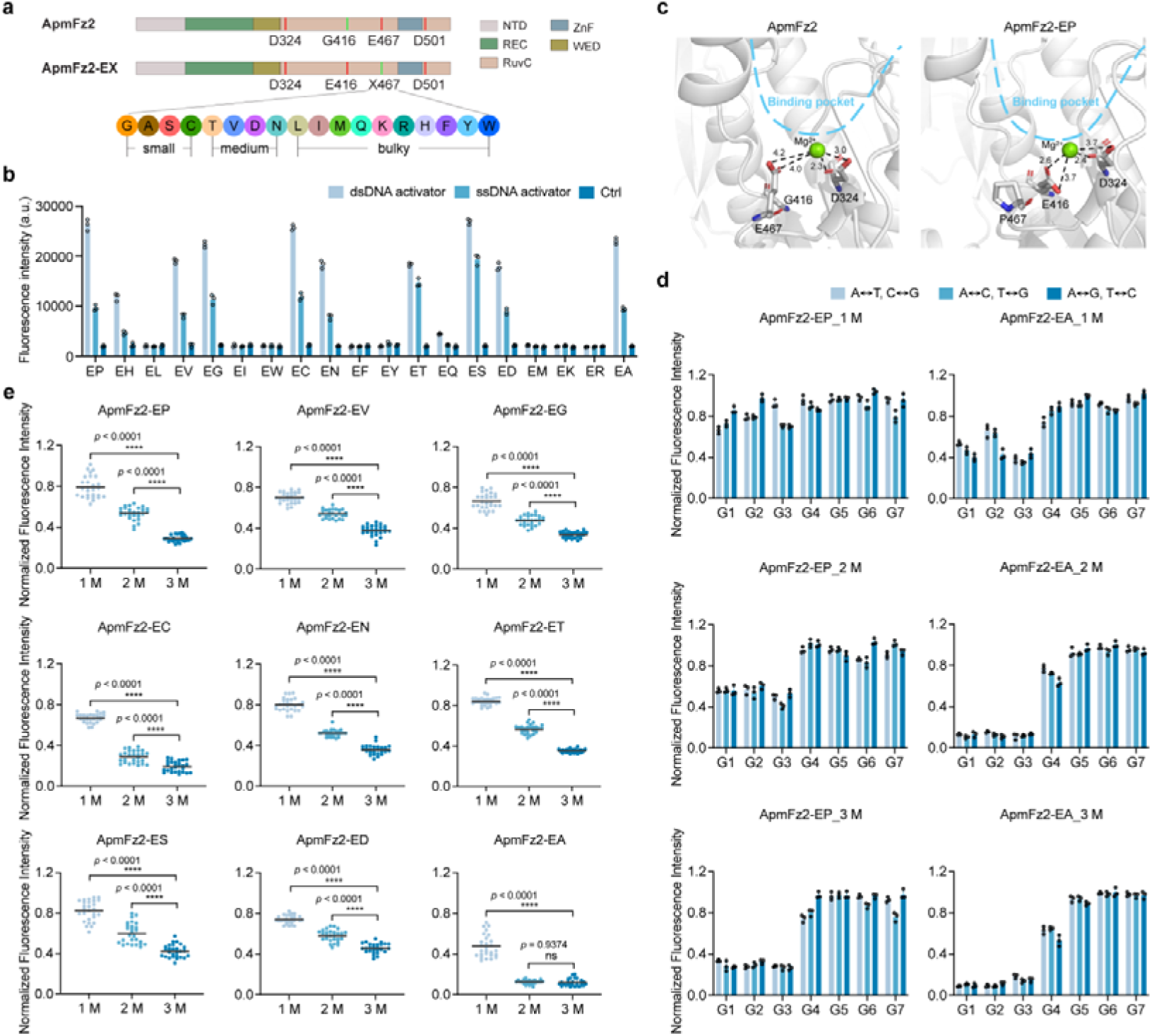
Residue 467 substitutions modulate ApmFz2 *trans*-cleavage activity and mismatch tolerance. **a**, Domain architectures of ApmFz2 and ApmFz2-EX variants. X indicates the amino acid substituted at position 467. Substitutions are grouped by side-chain size as small, medium, or bulky. **b**, *Trans*-cleavage activity of ApmFz2-EX variants in the presence of dsDNA or ssDNA activators. Ctrl, no-template control. Data are shown as mean ± s.d. (*n* = 3 technical replicates). **c**, Structural comparison of the catalytic centers of wild-type ApmFz2 (left) and ApmFz2-EP (right) using representative AlphaFold3 models. Distances between the Mg^2+^ ion and the side-chain carboxyl oxygen atoms of E416 or D324 are indicated by dashed lines and labeled in angstroms (Å). **d**, Normalized fluorescence signals of ApmFz2-EP and ApmFz2-EA against targets containing different mismatch positions and mismatch numbers. 1M, 2M, and 3M indicate targets containing one, two, or three mismatches, respectively. Data are shown as mean ± s.d. (*n* = 3 technical replicates). **e**, Average normalized fluorescence signals within the G1-G3 target region across mismatch-number categories for *trans*-cleavage-active ApmFz2 variants. 1M, 2M, and 3M indicate targets containing one, two, or three mismatches, respectively. Statistical significance was assessed by one-way analysis of variance (ANOVA). \*\*\*\**p* < 0.0001; ns, not significant. ApmFz2 variant proteins were purified as RNP complexes.

All *trans*-cleavage-active variants generated substantially stronger fluorescence signals after activation by dsDNA targets than by ssDNA targets (Fig. 3b). Notably, ApmFz2-ES and ApmFz2-ET showed only modest differences in fluorescence intensity between dsDNA and ssDNA activators. These results suggest that residue 467 modulates not only overall *trans*-cleavage activity but also the relative response of ApmFz2 variants to different nucleic acid targets.

Although the molecular basis of *trans*-cleavage activation remains incompletely defined, these observations led us to hypothesize that *cis*– and *trans*-cleavage activities use the same catalytic center but involve distinct conformational states that differentially affect substrate recognition and cleavage. To explore this possibility, we used AlphaFold3^28^ to predict structural models of all experimentally characterized ApmFz2 double variants in complex with the catalytic Mg^2+^ ion. All predicted models showed high confidence scores (pTM > 0.8 and ipTM > 0.8), supporting their use for comparable structural analysis (Extended Data Fig. 5). In the ApmFz2-EP model, the reconstructed G416E residue and the native catalytic residue D324 jointly coordinated the Mg^2+^ ion (Fig. 3c, right), supporting the functional role of the introduced glutamate in restoring catalytic geometry. The E467P substitution also removed steric bulk (Fig. 3c, right) that would otherwise be occupied by the native Glu467 side chain in wild-type ApmFz2 (Fig. 3c, left), creating space for the engineered E416 residue to adopt an extended, catalytically competent conformation. Consistent with this model, all *trans*-cleavage-active variants containing small-to-medium substitutions at position 467 adopted similar predicted active-site conformations (Extended Data Fig. 6).

In contrast, variants harboring long side-chain substitutions, such as arginine, lysine, or methionine at position 467, adopted extended conformations resembling that of native Glu467 (Extended Data Fig. 7). These side chains occupied a substantial portion of the active-site cavity and may interfere with nucleic acid binding or substrate positioning required for efficient *trans*-cleavage. Bulky aromatic substitutions, including phenylalanine, tyrosine, and tryptophan, adopted inward– or outward-facing conformations distinct from the native glutamate (Extended Data Fig. 7). Nevertheless, their large side-chain volumes likely impose steric constraints that limit substrate accommodation within the active site. Together, these AlphaFold3 models provide a plausible structural explanation for the activity differences observed among ApmFz2 double variants and suggest that side-chain volume at position 467 regulates *trans*-cleavage activation by modulating spatial accommodation within the catalytic pocket.

We next assessed how mismatch position and mismatch number affect the mismatch tolerance of *trans*-cleavage–active ApmFz2 variants. A 21-bp target region was divided into seven 3-bp groups (G1-G7), and one, two, or three mismatches were introduced within each group (Extended Data Fig. 8). These variants did not show an obvious preference for specific mismatch types, indicating that mismatch identity has limited influence on target recognition (Fig. 3d and Extended Data Fig. 9). For ApmFz2-EP, EV, EG, EN, ET, and ES, fluorescence signals gradually decreased as the number of mismatches increased within the G1-G3 region. In contrast, multiple consecutive mismatches in the TAM-distal G5-G7 region had minimal effect on fluorescence, with signals remaining close to wild-type levels (Fig. 3d,e and Extended Data Fig. 9). These results indicate a mismatch-sensitive, seed-like region spanning the TAM-proximal 9 nucleotides of the ωRNA. Compared with previously reported Cas12a or TnpB systems^15,29^, these ApmFz2 variants appeared to exhibit relatively high mismatch tolerance.

ApmFz2-EA and ApmFz2-EC showed substantially higher sensitivity to mismatches within the G1-G3 region. Among the tested variants, ApmFz2-EA exhibited the highest target specificity. Even a single mismatch caused a pronounced reduction in fluorescence signal, whereas two or three consecutive mismatches nearly abolished detectable activity (Fig. 3d,e and Extended Data Fig. 9). These results indicate that ApmFz2-EA has stricter target-recognition requirements and markedly reduced mismatch tolerance. Given this distinct specificity profile, we further examined its reporter cleavage preference under different target conditions. ApmFz2-EA showed a reporter preference similar to that of ApmFz2-EP (Extended Data Fig. 10).

Together, the distinct mismatch-tolerance profiles of these ApmFz2 variants suggest that they may be suited to different diagnostic applications. ApmFz2-EP, EV, EG, EN, ET, and ES, which exhibit broad mismatch tolerance, may be advantageous for broad-spectrum detection of highly homologous or multi-subtype pathogens. In contrast, the high mismatch sensitivity of ApmFz2-EA may support more stringent discrimination of single-nucleotide variations (SNV), making it potentially useful for SNV genotyping and subtype-specific detection.

### ApmFz2-EP enables broad detection across genetically diverse target subtypes

By integrating recombinase-aided amplification (RAA) with ApmFz2-EP *trans*-cleavage, we established FINDER, a Fanzor-based Integrated Nucleic acid DEtection platform (Fig. 4a). We first evaluated FINDER sensitivity by targeting the *L1* genes of human papillomavirus (HPV) subtypes 16 and 18 using subtype-specific ωRNAs. The assay achieved limits of detection as low as ∼1.6 copies/μL for HPV16 and ∼5 copies/μL for HPV18 (Fig. 4b). To assess clinical applicability, we analyzed HPV16/18 infection in 27 liquid-based thin-layer cytology (TCT) samples using quantitative PCR (qPCR) as the reference standard. FINDER showed complete concordance with qPCR detection (Fig. 4c,d and Extended data Fig. 11). The assay also produced fluorescence signals that were readily distinguishable under blue-light illumination, supporting its potential for visual nucleic acid detection.

**Fig. 4.**
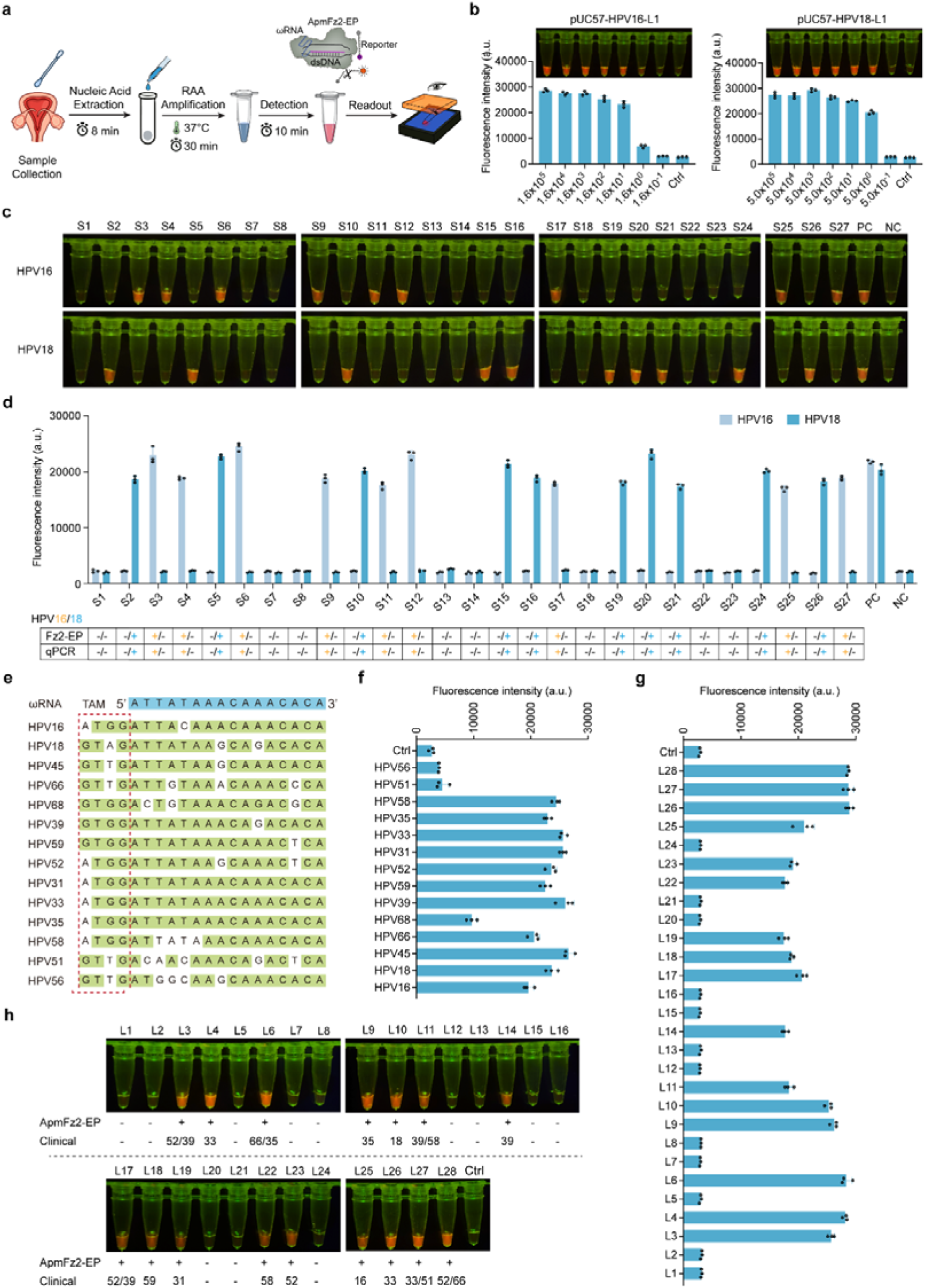
ApmFz2-EP-mediated FINDER enables sensitive HPV detection and broad recognition of high-risk HPV subtypes. **a**, Schematic of the FINDER workflow, including sample collection, nucleic acid extraction, RAA amplification, ApmFz2-EP-mediated reporter cleavage, and fluorescence readout. **b**, Analytical sensitivity of FINDER for HPV16 and HPV18 detection using plasmids containing the corresponding *L1* genes as templates. Ctrl, no-template control. Data are shown as mean ± s.d. (*n* = 3 technical replicates). **c,d**, Validation of FINDER-based HPV16 and HPV18 detection in clinical samples using blue-light visualization (c) and fluorescence intensity measurements (d). PC, positive control containing a pUC57-L1 plasmid; NC, no-template negative control; +, positive samples; −, negative samples. Data are shown as mean ± s.d. (*n* = 3 technical replicates). **e**, Sequence alignment of target regions from 14 high-risk human papillomavirus (HR-HPV) subtypes used for universal ωRNA design. The blue sequence indicates the universal ωRNA sequence, and red boxes denote the corresponding TAM sequences. **f**, Detection of 14 HR-HPV subtypes by FINDER using the universal ωRNA. Ctrl, no-template control. Data are shown as mean ± s.d. (*n* = 3 technical replicates). **g,h**, Validation of FINDER for broad HR-HPV subtype detection in clinical samples using fluorescence intensity measurements (g) and blue-light visualization (h). Ctrl, no-template control; +, positive samples; −, negative samples. Data are shown as mean ± s.d. (*n* = 3 technical replicates). ApmFz2-EP protein was purified using the pCold-TF expression system.

Because ApmFz2-EP showed broad mismatch tolerance and relaxed TAM requirements, we next tested whether a single ωRNA could detect multiple genetically diverse HPV subtypes. This strategy could reduce the need for multiple guide RNAs, which often require Cas12a-based assays to accommodate viral sequence diversity. We compared *L1* gene sequences from 14 high-risk HPV subtypes (52, 66, 39, 59, 56, 45, 68, 51, 35, 33, 16, 31, 18, and 58) and designed a universal ωRNA targeting a relatively conserved region compatible with variable TAM contexts (Fig. 4e). Within this design, HPV51 and HPV56 did not generate detectable fluorescence, and HPV68 produced only a weak response. In contrast, the remaining 11 HPV subtypes efficiently activated ApmFz2-EP *trans*-cleavage and produced strong fluorescence signals despite sequence variation within the target region (Fig. 4f). We further evaluated this broad-detection strategy in 28 clinical specimens infected with different high-risk HPV types. FINDER generated stable fluorescence signals across HPV-positive samples of different subtypes and showed high concordance with clinical diagnostic assays (Fig. 4g,h).

We also tested whether *trans*-cleavage-active ApmFz2 variants could support a one-pot amplification-detection format. Among the tested variants, only ApmFz2-ET and ApmFz2-ES enabled one-pot detection (Extended data Fig. 12a,b), indicating that reaction optimization may be needed for fully integrated formats. To evaluate the clinical applicability of the one-pot format, we used the ApmFz2-ES variant to analyze 17 porcine blood samples for African swine fever virus (ASFV) detection. The assay showed complete concordance with quantitative PCR (qPCR), correctly identifying all ASFV-positive and ASFV-negative samples (Extended Data Fig. 12c-e). Together, these results show that the ApmFz2-EP-based FINDER platform supports sensitive detection of individual HPV subtypes and broad recognition of genetically diverse high-risk HPV targets.

### ApmFz2-EA enables single-nucleotide variant discrimination

We next asked whether the mismatch-sensitive ApmFz2-EA variant could be used for SNV discrimination. As a model target, we selected the sheep FecB locus, a prolificacy-associated *BMPR1B* variant caused by an A746G substitution. By combining RAA with ApmFz2-EA *trans*-cleavage, we developed a FINDER-based SNV genotyping platform assay for this locus (Fig. 5a). Allele-specific ωRNAs were designed to distinguish the AA and GG genotypes, and the TAM sequence was introduced into the RAA amplicon through primer design (Fig. 5b). Using genomic DNA (gDNA) extracted from sheep blood, the assay detected as little as 0.8 pg of input gDNA (Extended data Fig. 13).

**Fig. 5.**
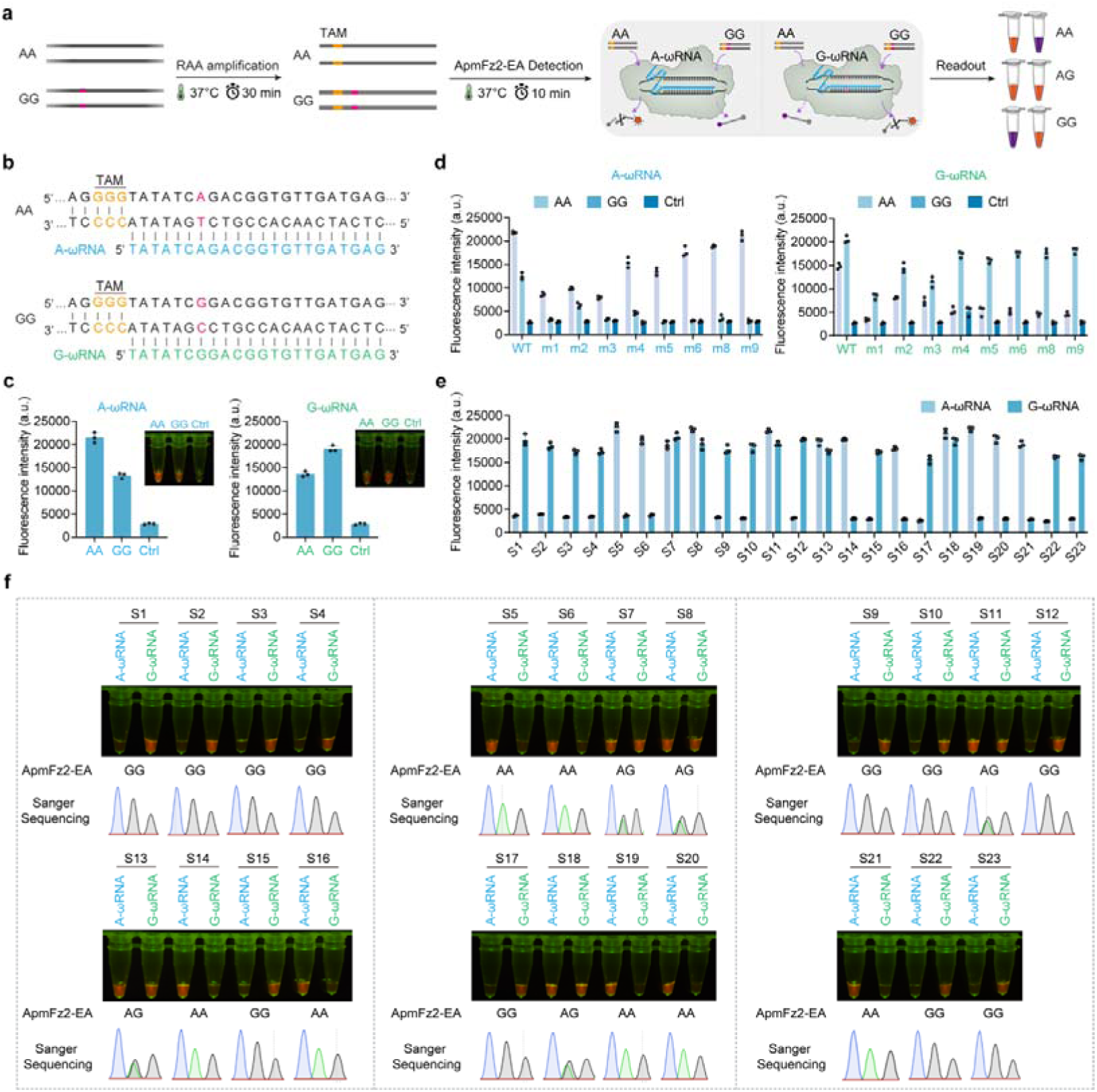
ApmFz2-EA-mediated FINDER enables single-nucleotide variant discrimination. **a**, Schematic of the FINDER workflow for FecB genotyping, including RAA amplification, ApmFz2-EA-mediated reporter cleavage, and genotype readout. **b**, Design of allele-specific A-ωRNA and G-ωRNA for discrimination of the AA and GG genotypes at the FecB locus. The TAM sequence is indicated in orange, and the genotype-defining nucleotide is highlighted in pink. **c**, Initial evaluation of allele-specific A-ωRNA and G-ωRNA for FecB genotyping. Ctrl, no-template control. Data are shown as mean ± s.d. (*n* = 3 technical replicates). **d**, Optimization of engineered secondary mismatches within the ωRNA-target duplex to improve allelic discrimination. WT indicates the original allele-specific ωRNA without an engineered secondary mismatch; m1-m9 indicate engineered mismatches at positions 1-9. Ctrl, no-template control. Data are shown as mean ± s.d. (*n* = 3 technical replicates). **e,f**, Validation of ApmFz2-EA-mediated FINDER for FecB genotyping in 23 sheep blood samples using fluorescence intensity measurements (e) and blue-light visualization with corresponding Sanger sequencing chromatograms (f). S1-S23 denote individual sheep blood samples. Data are shown as mean ± s.d. (*n* = 3 technical replicates). ApmFz2-EP protein was purified using the pCold-TF expression system.

We then tested whether the allele-specific ωRNAs could reliably distinguish the two genotypes. Matched ωRNA-target pairs produced higher fluorescence signals than mismatched pairs, but the signal distributions still overlapped substantially between AA and GG samples (Fig. 5c). This overlap limited genotype discrimination and indicated that allele-specific ωRNA design alone was insufficient.

To improve specificity, we introduced a deliberate secondary mismatch at positions 1 to 9 within the ωRNA-target duplex (Fig. 5d). Secondary mismatches near the 5’ end of the target region, especially positions 1-3, increased discrimination but also sharply reduced signal from matched targets. This loss of signal limited their practical utility. In contrast, mismatches at positions 6, 8, or 9 retained robust activation by matched targets while suppressing fluorescence from mismatched alleles. Among these designs, the position-9 mismatch provided the best balance between signal intensity and genotype separation. The same pattern was observed with the G-ωRNA design, in which mismatches at positions 6, 8, or 9 reduced background signal from the AA genotype while preserving strong signal from the matched GG genotype (Fig. 5d). Based on these results, we selected the position-9 engineered mismatch for subsequent genotyping assays.

Finally, we evaluated the optimized assay using 23 sheep blood samples. ApmFz2-EA-based genotype calls were fully concordant with Sanger sequencing results (Fig. 5e,f), support the accuracy of the assay in the tested sample set. These findings show that ApmFz2-EA, when paired with an engineered mismatch strategy, can support specific SNV discrimination within the FINDER platform.

## Discussion

RNA-guided nucleases have become important tools for nucleic acid detection because target binding can activate collateral cleavage of reporter substrates. This principle has been used with Cas12, Cas13, TnpB, and, more recently Cas9-based systems^6,15,30^, although each system has distinct biochemical and practical constraints. In this study, we show that Fanzors, a class of eukaryotic RNA-guided nucleases, can be engineered to acquire *trans*-cleavage activity. Rewiring the catalytic center of ApmFz2 activated collateral cleavage while substantially reducing canonical *cis*-cleavage. When paired with isothermal amplification, this activity supported a Fanzor-based nucleic acid detection platform, FINDER, for pathogen detection and SNV genotyping.

The central finding of this work is that unusual Fanzor catalytic architecture helps determine nuclease behavior. In Cas12 and TnpB systems, *trans*-cleavage is linked to an activated RuvC state that permits nonspecific cleavage after target recognition ^6,10,12,25,31^. Fanzors differ from canonical TnpB proteins in that the catalytic glutamate within the RuvC-II motif is displaced toward the C terminus, producing a noncanonical “D-altE-D” arrangement^10,17,18,32^. By restoring a TnpB-like “D-E-D” configuration in ApmFz2, we activated *trans*-cleavage activity that is weak or undetectable in the wild-type enzyme. This result indicates that the catalytic-center geometry of ApmFz2 is a major determinant of its cleavage mode. It also suggests that Fanzors retain a latent capacity for collateral cleavage that may have been attenuated during evolution through remodeling of the catalytic core.

Our mutational analysis further showed that *trans*-cleavage activation depends not only on the presence of the reconstructed glutamate but also on the local steric environment around the active site. Variants with small-to-medium side chains at position 467 retained *trans*-cleavage activity, whereas most bulky substitution greatly reduced or abolished activity (Fig. 3b). AlphaFold3 structural models were consistent with this pattern, suggesting that smaller residues at position 467 create space that allows the engineered E416 residue to adopt a catalytically favorable conformation. This observation is conceptually similar to the “open active-site” model proposed for Cas12a, in which local rearrangements around catalytic residues regulate substrate access and active-site exposure^25,33–36^. Although the activated ApmFz2 structure has not yet been resolved experimentally, biochemical and modeling data support a model in which side-chain volume near the catalytic pocket helps determine whether ApmFz2 can accommodate nonspecific reporter substrates after target recognition.

Distinct substitutions at position 467 also changed target-recognition behavior (Fig. 3e and Extended data Fig. 9). ApmFz2-EP and several related variants tolerated mismatches across much of the target sequence, with the strongest mismatch sensitivity concentrated in the TAM-proximal region. This broad tolerance allowed a single ωRNA to detect multiple high-risk HPV subtypes, including targets with sequence variation in the recognition region. ApmFz2-EA also required only seven nucleotides of guide-target complementarity to activate *trans*-cleavage, which is substantially shorter than the activation thresholds reported for Cas12a (≥14 nt) and several TnpB systems (≥12 nt)^15,26,37^. This unusually short requirement may reflect a target-recognition mechanism that differs from those of Cas12 and TnpB, although structural and biochemical studies will be needed to define that mechanism.

Engineered Fanzors may also complement existing Cas12-, Cas13-, and TnpB-based diagnostic systems. First, ApmFz2-EP can be activated by both dsDNA and ssRNA activators and shows different reporter preferences depending on the activating nucleic acid. DNA-triggered activation supported cleavage of poly(T)-, poly(C)-, and poly(A)-containing ssDNA reporters, whereas RNA-triggered activation preferentially induced cleavage of poly(C) reporters (Fig. 2d). This separation between activator type and reporter preference may be useful for multiplex assay design. For example, combining engineered Fanzors with Cas13 systems, which cleave ssRNA reporters^2,5,38–40^, could allow simultaneous detection of multiple DNA and RNA targets in a single reaction. Second, ApmFz2-EP appears to have more flexible target-adjacent sequence requirements than many Cas12 effectors, which often depend on T-rich PAM sequences and therefore constrained target selection^8,41,42^ (Fig. 2b). This different targeting behavior could expand the range of accessible diagnostic sites, especially in regions that are poorly suited to Cas12-based detection. Third, ApmFz2-EP can be activated by as few as seven nucleotides of guide-target complementarity (Fig. 2c). This short activation requirement may be useful for detecting fragmented nucleic acid, including cell-free DNA and environmental DNA, although those applications will need to be tested directly. Finally, the reduction in *cis*-cleavage activity may help preserve target molecules and extend the lifetime of the activated target-bound complex, thereby supporting sustained reporter cleavage and stronger signal amplification.

Despite these advances, several limitations should be considered. First, the structural basis of *trans*-cleavage activation remains unresolved. Although the AlphaFold3 models support a steric mechanism, they do not replace experimental structures. Cryo-electron microscopy or other structural approaches will be needed to define the conformational changes associated with activation of the *trans*-cleavage state. Second, although engineered ApmFz2 variants showed robust *trans*-cleavage activity, their catalytic efficiency remains lower than that of some extensively optimized Cas12a-based diagnostic systems^43–45^. Additional improvement through directed evolution, structure-guided engineering^46,47^, and guide RNA optimization may be needed to improve catalytic efficiency and diagnostic sensitivity. Third, this study focused mainly on ApmFz2 as a representative Fanzor effector. It remains unclear whether a similar catalytic-center rewiring strategy will work across the broader Fanzor family, so additional homologs will need to be tested. Finally, despite their compact size, engineered Fanzors currently appear less compatible with one-pot amplification-detection workflows than established one-pot detection platforms^15,48^. Further optimization of reaction conditions and effector properties may be required before fully integrated one-pot Fanzor-based assays are practical.

In summary, this study shows that catalytic-center rewiring can give the eukaryotic RNA-guided nuclease ApmFz2 robust *trans*-cleavage activity. It also identifies the amino acid composition near the catalytic site as a key determinant of both catalytic efficiency and target-recognition stringency. These findings clarify how Fanzor catalytic architecture shapes nuclease function and provide design principles for engineering compact RNA-guided nucleases for nucleic acid detection.

## Methods

### Plasmid and double-stranded DNA target preparation

The coding sequence of ApmFz2 was codon-optimized for expression in *Escherichia coli*, synthesized by GenScript Biotech (Nanjing, China), and cloned into the pET-28a vector. For protein expression, ApmFz2 and its variants were subcloned into the appropriate expression vectors, including pCold-TF (TaKaRa) and a modified pET-28a vector containing an N-terminal Twin-Strep tag for ribonucleoprotein (RNP) purification. ApmFz2 variants were generated by site-directed mutagenesis using the QuickMutation™ Site-Directed Mutagenesis Kit (Beyotime Biotechnology, China) according to the manufacturer’s instructions. All mutations were confirmed by Sanger sequencing before subsequent experiments, and the corresponding amino acid sequences of all ApmFz2 variants are provided in Supplementary Table S1.

All ωRNA expression constructs were generated by cloning synthesized ωRNA coding sequences into the pCDFDuet-1 vector (Novagen). ASFV-*p72*, CSFV-*E2*, HPV16-*L1*, and HPV18-*L1* gene fragments were synthesized and cloned into pUC57 by GenScript Biotech to generate pUC57-ASFV-p72, pUC57-CSFV-E2, pUC57-HPV16-L1 and pUC57-HPV18-L1, respectively. These plasmids were used as templates for PCR amplification, analytical sensitivity evaluation, and one-pot detection assays.

Double-stranded DNA (dsDNA) targets used in this study, including 30 TAM variants, dsDNA activators with different target lengths, and targets containing mismatches at different positions, of different types, or in different numbers, were generated by PCR amplification using specifically designed primers. PCR products were analyzed by agarose gel electrophoresis, and fragments of the expected size were purified using either a DNA Gel Extraction Kit or a PCR Purification Kit (Omega). DNA concentration and purity were measured using a NanoDrop 2000 spectrophotometer (Thermo Fisher Scientific). Purified dsDNA targets were aliquoted and stored at –20 °C until use.

### Recombinant protein expression and purification

The coding sequences of ApmFz2 and its variants were inserted into the pCold-TF expression vector, which encodes an N-terminal His_6_-Trigger Factor (TF) solubility tag followed by a human rhinovirus (HRV) 3C protease cleavage site. The resulting plasmids were transformed into *Escherichia coli* BL21(DE3) cells for recombinant protein expression. Single colonies were used to inoculate LB medium containing 100 μg mL^-1^ ampicillin, and cultures were grown overnight at 37 °C with shaking at 200 rpm. Overnight cultures were diluted 1:100 into 3 L of fresh LB medium supplemented with ampicillin and grown at 37 °C until the OD_600_ reached 0.6 to 0.8. The cultures were chilled on ice for 30 min, induced with 0.5 mM IPTG, and incubated at 16 °C for 16 h to allow protein expression.

Cells were collected by centrifugation at 9,000 × g for 15 min at 4 °C and resuspended in lysis buffer composed of 50 mM HEPES (pH 7.5), 300 mM NaCl, 10 mM imidazole, 5% (v/v) glycerol, 1 mM DTT, and 0.1× cOmplete™ EDTA-free protease inhibitor cocktail (Millipore). The cell suspension was lysed by sonication on ice, and insoluble material was removed by centrifugation at 12,000 × g for 1 h at 4 °C. The clarified lysate was incubated for 2 h at 4 °C with Ni-NTA agarose resin (Cytiva) that had been equilibrated with lysis buffer, and the mixture was then transferred to a gravity-flow column. The resin was washed with buffer containing 20 to 50 mM imidazole, and bound proteins were eluted with 80 to 300 mM imidazole in 10 mM Tris-HCl (pH 7.5) and 500 mM NaCl. Eluted fractions were examined by SDS-PAGE, and fractions containing the target protein were pooled and concentrated using Amicon Ultra centrifugal filter units (Merck).

The concentrated protein sample was further purified by size-exclusion chromatography on a Superdex 200 Increase 10/300 GL column (Cytiva) equilibrated in 10 mM Tris-HCl (pH 7.8), 300 mM NaCl, and 0.5 mM DTT. Peak fractions were analyzed by SDS-PAGE, pooled, and treated with HRV 3C to remove the N-terminal tag. For cleavage, purified fusion proteins were incubated with HRV 3C protease (TaKaRa) at a protease-to-substrate mass ratio of 1:60 for 15 h at 4 °C. The digestion mixture was then incubated with TALON Metal Affinity Resin (TaKaRa) for 2 h at 4 °C with gentle end-over-end rotation. His_6_-TF, uncleaved fusion protein, and His-tagged HRV 3C protease were retained on the resin, whereas tag-free ApmFz2 remained in the unbound fraction. After centrifugation at 700 × g for 5 min, the supernatant containing purified tag-free ApmFz2 was collected.

Purified proteins were dialyzed overnight at 4 °C against storage buffer containing 20 mM sodium acetate (pH 6.0), 500 mM NaCl, 0.1 mM EDTA, 0.1 mM tris(2-carboxyethyl)phosphine (TCEP), and 50% (v/v) glycerol. Protein concentrations were measured using a BCA Protein Assay Kit (Beyotime Biotechnology, China). The purified proteins were aliquoted and stored at –80 °C until use.

### Ribonucleoprotein purification

To purify ApmFz2 ribonucleoprotein (RNP) complexes, the coding sequences of ApmFz2 and its variants were modified to encode an N-terminal Twin-Strep tag and cloned into the *Nco* I and *Xho* I sites of the pET-28a expression vector. The corresponding ωRNA sequences were cloned individually into the pCDFDuet-1 vector. Protein-expression and ωRNA-expression plasmids were co-transformed into *Escherichia coli* BL21(DE3) competent cells to allow intracellular assembly of ApmFz2 RNP complexes.

Single colonies were used to inoculate Terrific Broth (TB) medium supplemented with 50 μg mL^-1^ kanamycin and 50 μg mL^-1^ streptomycin, and cultures were grown overnight at 37 °C with shaking at 200 rpm. Overnight culture was transferred into 2 L of fresh TB medium containing the same antibiotics and grown at 37 °C until the OD_600_ reached 0.6 to 0.8. Cultures were then chilled at 4 °C for 30 min, induced with 0.5 mM IPTG, and incubated for an additional 16 h at 16 °C.

Cells were harvested by centrifugation at 9,000 × g for 15 min at 4 °C and resuspended in lysis buffer containing 50 mM HEPES (pH 7.5), 300 mM NaCl, 5% (v/v) glycerol, 1 mM DTT, and 0.1× cOmplete™ EDTA-free protease inhibitor cocktail (Millipore). The cell suspension was lysed by sonication on ice, and insoluble debris was removed by centrifugation at 12,000 × g for 1 h at 4 °C. The clarified supernatant was incubated with pre-equilibrated Strep-Tactin^®^XT 4Flow^®^ high capacity resin (IBA Lifesciences) for 3 h at 4°C with gentle rotation. The resin was then transferred to a gravity-flow column and washed extensively with Buffer W (IBA Lifesciences). Bound RNP complexes were eluted with Buffer BXT (IBA Lifesciences), and elution fractions were collected separately.

Aliquots of each elution fraction (20 μL) were analyzed by SDS-PAGE. Fractions containing highly pure ApmFz2 RNP complexes were pooled and concentrated using Amicon Ultra centrifugal filter units (Merck). The concentrated RNP complexes were dialyzed overnight at 4 °C, 500 mM NaCl, 0.1 mM EDTA, 0.1 mM TCEP, and 50% (v/v) glycerol. RNP concentrations were measured using a BCA Protein Assay Kit (Beyotime Biotechnology), and samples were aliquoted and stored at –80 °C until use.

### *In vitro* transcription

All ωRNAs and target RNAs used in this study were prepared by *in vitro* transcription (IVT) using T7 RNA polymerase. DNA templates were designed to contain a T7 promoter upstream of the target RNA sequence, synthesized by Tsingke Biotechnology, and amplified by PCR before transcription. IVT reactions were performed using the HiScribe™ T7 High Yield RNA Synthesis Kit (New England Biolabs) according to the manufacturer’s instructions. After transcription, residual template DNA was digested with DNase I treatment at 37 °C for 15 min. RNA products were purified with the Monarch^®^ RNA Cleanup Kit (New England Biolabs) following the manufacturer’s protocol and quantified using a NanoDrop 2000 spectrophotometer. Purified ωRNAs and target RNAs were diluted to the required concentrations in RNase-free water, aliquoted to avoid repeated freeze-thaw cycles, and stored at –80 °C. The sequences of all ωRNAs used in this study are listed in Supplementary Table S2.

### *cis*-cleavage assay

Cy5– and FAM-labeled oligonucleotides were synthesized by GenScript Biotech and used as primers to amplify target-containing DNA fragments for *cis*-cleavage assays. These PCR products served as fluorescently labeled dsDNA substrates. Cleavage reactions were assembled in a total volume of 20 μL and contained 1× Fz Buffer 4.0 [10 mM KCl, 40 mM Tris-HCl, 20 mM MgCl_2_, 0.001% Triton X-100, 100 μg mL^-1^ BSA, 2mM DTT, pH 8.5], 500 nM ApmFz2 or variant protein, 300 nM corresponding ωRNA, and 120 ng fluorescently labeled dsDNA substrate. Unless otherwise indicated, reactions were incubated at 37 °C for the specified time periods. Reactions were stopped by adding Proteinase K (New England Biolabs), followed by incubation at 60 °C for 5 min. An equal volume of 100% formamide (Thermo Fisher Scientific) was then added, and samples were denatured at 95 °C for 5 min. Cleavage products were separated on 12% TBE-urea polyacrylamide gels (Thermo Fisher Scientific) and visualized using a fluorescence imaging system (Sinsage, China). FAM– and Cy5-labeled products were detected in the green and red channels, respectively.

### ApmFz2 *trans*-cleavage assay

*Trans*-cleavage assays were generally assembled in 20 μL reactions containing reaction buffer, ApmFz2 or the indicated variants, ωRNA, a target nucleic acid activator, and a fluorescent reporter substrate. Unless otherwise stated, standard reactions contained 500 nM ApmFz2 or variant protein, 300 nM ωRNA, 50 nM target DNA activator or 1 μL RAA amplicon, and 250 nM reporter substrate in 1× reaction buffer. For experiments using preassembled RNP complexes, reactions contained 900 nM purified RNP, 50 nM target DNA activator or 1 μL RAA amplicon, and 250 nM reporter substrate in 1× reaction buffer. All modified reporter sequences used in this study are listed in Supplementary Table S4.

For gel-based *trans*-cleavage assays, reactions were incubated at 37 °C for the indicated time, then terminated by adding Proteinase K and incubated at 60 °C for 5 min. An equal volume of 100% formamide was added, and samples were denatured at 95 °C for 5 min. Reaction products were separated on 15% denaturing TBE-urea polyacrylamide gels (Thermo Fisher Scientific) and visualized using a fluorescence imaging system with the appropriate detection channels.

For fluorescence-based *trans*-cleavage assays, reactions were assembled in PCR tubes and included a 250 nM ssDNA fluorescent reporter labeled with ROX and BHQ2 (5′-ROX/GTATCCAGTGCG/BHQ2-3′). Reactions were incubated at 37 °C for the indicated time and terminated by heating at 98 °C for 2 min. Products were diluted with 80 μL nuclease-free water and transferred to black 96-well plates for fluorescence measurement. Fluorescence was measured using a Synergy H1 multimode microplate reader (BioTek) with excitation and emission wavelengths of 576 nm and 601 nm, respectively. For endpoint assays, fluorescence was recorded immediately after samples were transferred to the plates. For kinetic assays, reactions were assembled directly in 96-well plates, incubated at 37 °C, and monitored every 1 min.

### ApmFz2 *trans*-cleavage reaction condition optimization

To optimize ApmFz2-EP *trans*-cleavage conditions, key buffer components were evaluated systematically using fluorescence-based *trans*-cleavage assays. Unless otherwise noted, reactions were performed under identical conditions, with only the tested parameter varied. For pH optimization, reactions were assembled in buffers ranging from pH 7.0 to 9.0. The buffer that produced the highest activity was designated Fz Buffer 1.0. Mg^2+^ concentration was then optimized by varying MgCl_2_ from 5 to 30 mM, using Fz Buffer 1.0 as the base buffer, to generate Fz Buffer 2.0. Reducing conditions were evaluated next by testing DTT concentrations from 0 to 10 mM in Fz Buffer 2.0, yielding Fz Buffer 3.0. Finally, monovalent salt conditions were optimized by varying KCl from 10 to 50 mM in Fz Buffer 3.0. The resulting optimized formulation was designated Fz Buffer 4.0 and used for all subsequent experiments.

### Recombinase-aided amplification

Recombinase-aided amplification (RAA) was performed using a commercial RAA kit (Qitian Bio, China), following the manufacturer’s instructions with minor modifications. For each standard reaction setup, 25 μL Buffer V, one lyophilized reaction pellet, and 5 μL magnesium acetate [Mg(OAc)_2_] were combined to reconstitute the amplification mixture. The reconstituted mixture was then divided equally into individual amplification reactions. Each 25 μL reaction contained 1 μL of 10 μM forward primer, 1 μL of 10 μM reverse primer, and either 4 μL of crude DNA extract or 1 μL of plasmid DNA template. Reactions were incubated at 37 °C for 30 min in a metal heating block and then heat-inactivated at 80 °C for 5 min. The resulting amplicons were used directly in downstream ApmFz2-based detection assays. All primer and oligonucleotide sequences used in this study are listed in Supplementary Table S3.

### One-pot ApmFz2-mediated nucleic acid detection

The one-pot ApmFz2 detection assay combined RAA amplification and ApmFz2-mediated *trans*-cleavage in a single isothermal reaction. Reactions were assembled using an RAA kit and included 25 μL Buffer V, 5 μL Mg(OAc)_2_, and one lyophilized reaction pellet. For each assay, half of the standard RAA reaction mixture was supplemented with 0.8 μL each of 10 μM forward and reverse primers, 700 nM purified ApmFz2 variant RNP, 250 nM ssDNA fluorescent reporter (5′-ROX/GTATCCAGTGCG/BHQ2-3′), and the corresponding nucleic acid template. The final reaction volume was adjusted to 25 μL. Reactions were incubated at 37 °C for 40 min in a metal heating block to allow target amplification and reporter cleavage to occur in the same tube, then heat-inactivated at 80 °C for 5 min. For endpoint visualization, reaction tubes were illuminated directly with a blue-light transilluminator, and fluorescence signals were captured with a smartphone camera. For quantitative analysis, reaction products were diluted with nuclease-free water to a final volume of 100 μL and transferred to black 96-well plates. Fluorescence was measured using a Synergy H1 microplate reader with excitation and emission wavelengths of 576 nm and 601 nm, respectively.

### Sample collection and DNA extraction

Sheep blood samples were collected for FecB genotyping. Clinical samples used for African swine fever virus (ASFV) detection were obtained from porcine blood, whereas samples used for human papillomavirus (HPV) detection consisted of ThinPrep cytology test (TCT) preservation fluids provided by Peking University Shenzhen Hospital. Crude genomic DNA was extracted using QuickExtract™ DNA Extraction Solution (Lucigen) according to the manufacturer’s protocol, with minor modifications. Briefly, 50 μL of each sample was mixed with 70 μL of QuickExtract™ solution and incubated at 65 °C for 8 min, followed by 98 °C for 2 min to complete lysis and inactivate enzymes. The resulting lysate was used directly as the template for downstream amplification and detection without further purification. Extracted DNA was stored at −20 °C until use.

### Quantitative PCR assays

Quantitative PCR (qPCR) was performed on a LightCycler^®^ 480 Instrument II (Roche). HPV16 and HPV18 were detected using TaqMan probe-based qPCR assays with Luna^®^ Universal Probe qPCR Master Mix (New England Biolabs), following the manufacturer’s protocol. Each 20 μL reaction contained 10 μL of 2× Luna^®^ Universal Probe qPCR Master Mix, gene-specific primers and probes, and 2 μL of extracted DNA template. Reactions were run with an initial denaturation step at 95 °C for 3 min, followed by 40 cycles of 95 °C for 10 s and 60 °C for 30 s. Fluorescence was collected during the extension step. Cycle threshold (Ct) values were calculated automatically using LightCycler^®^ 480 software (Roche) with the default analysis settings. All samples were analyzed with positive and negative controls. Primer and probe sequences used for PCR are listed in Supplementary Table S3.

### Structure modeling of ApmFz2 variants

The cryo-EM structure of wild-type ApmFz2 bound to DNA and RNA molecules was obtained from the Protein Data Bank (PDB ID: 9B0L). Structural inspection in PyMOL showed that the N-terminal 52 amino acids are absent from the experimentally determined structure, likely because this region is flexible and lacks well-defined electron density. To improve the confidence and consistency of the AlphaFold3 predictions, all structural models were generated using an ApmFz2 sequence lacking the first 52 amino acids.

Structural models were predicted using a locally installed version of AlphaFold3 (version 3.0.0). Nineteen engineered double variants, including ApmFz2-EA, EC, ED, EF, EG, EH, EI, EK, EL, EM, EN, EP, EQ, ER, ES, ET, EV, EW, and EY, were modeled together with the wild-type ApmFz2-WT control. To improve reproducibility, AlphaFold3 was run using five random seeds (1, 2, 3, 4, and 5) for each variant, and five diffusion samples were generated per seed, resulting in 25 structural models for each sequence. Unless otherwise indicated, predictions were performed using the default AlphaFold3 parameters. For each variant, the top-ranked model identified by the AlphaFold3 internal ranking metric was selected for downstream structural analysis.

## Statistical analysis

All experiments were performed at least three times independently unless otherwise indicated. Data are presented as mean ± standard deviation (s.d.). Statistical analyses were conducted using GraphPad Prism 9.0. Differences between groups were evaluated by one-way analysis of variance (ANOVA). Statistical significance was defined as ns, not significant, and \*\*\*\**p* < 0.0001.

## Ethical statement

The use of human samples for HPV testing from a cervical cancer prevention cohort at Peking University Shenzhen Hospital (PUSH) was approved by the Ethics Committee of PUSH (IRB No. PUSH [2023]-016). ASFV samples were provided by the China Animal Health and Epidemiology Center. All animal-related experiments were approved under ethics approval number DWFL-2026-05.

### Acknowledgements

We thank Prof. Jianlin Han (from Yazhouwan National Laboratory) for providing sheep blood samples and Prof. Di Wu (from Peking University Shenzhen Hospital) for providing HPV clinical samples. We also thank the China Animal Health and Epidemiology Center for providing ASFV samples. We acknowledge the Advanced Research Computing (ARC) facility at the University of Michigan for providing computational resources and services that supported this work. This work was supported by the National Key Research and Development Program of China (2023YFF1001000), the Fundamental Research Funds for the Central Universities (2662025DKPY005), the Basic Research Project of Yazhouwan National Laboratory (2310SH01), the Agricultural Gene Editing Platform Technology and Breeding Program of Hubei (2024BBA001), and the China Agricultural Research System (CARS-35).

## Author contributions

C.Z. and S.Z. conceived and supervised the study. C.Z. and B.X. designed the experiments. B.X. performed all *trans*-cleavage and biochemical assays and purified proteins. X.H. performed structure modeling and analysis. C.Z., B.X., and X.H. analyzed the data and prepared the figures. C.Z. and B.X. wrote the original draft of the manuscript. C.Z., X.H., and S.Z. critically revised the manuscript. All authors reviewed and approved the final version.

