## Supplementary Figures for "Catalytic rewiring of RuvC-II catalytic site activates *trans*-cleavage in Fanzor2"

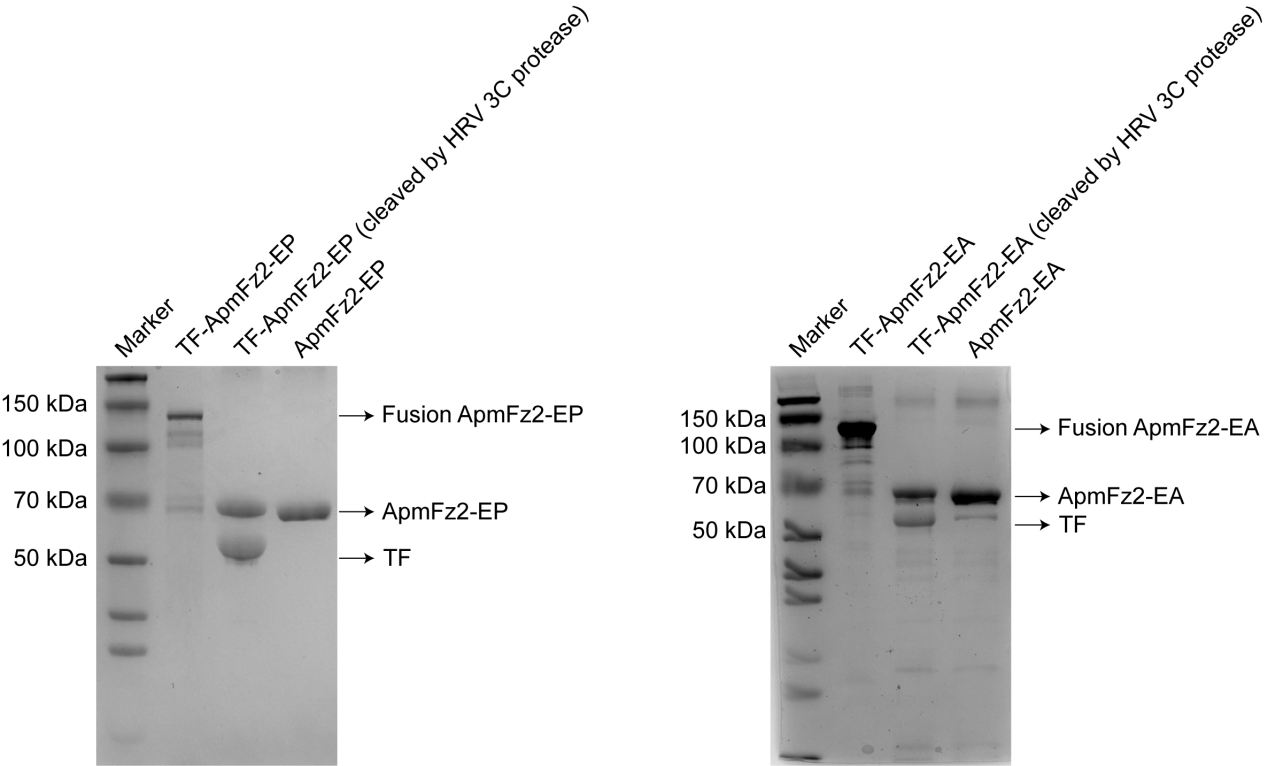

**Extended Data Fig. 1 SDS-PAGE analysis of ApmFz2-EP and ApmFz2-EA purification.** Coomassie Brilliant Blue-stained SDS-PAGE gels showing expression and purification of ApmFz2-EP and ApmFz2-EA proteins. Fusion proteins, HRV 3C-cleaved products, purified tag-free ApmFz2 variants, and the released Trigger Finger (TF) tag are indicated.

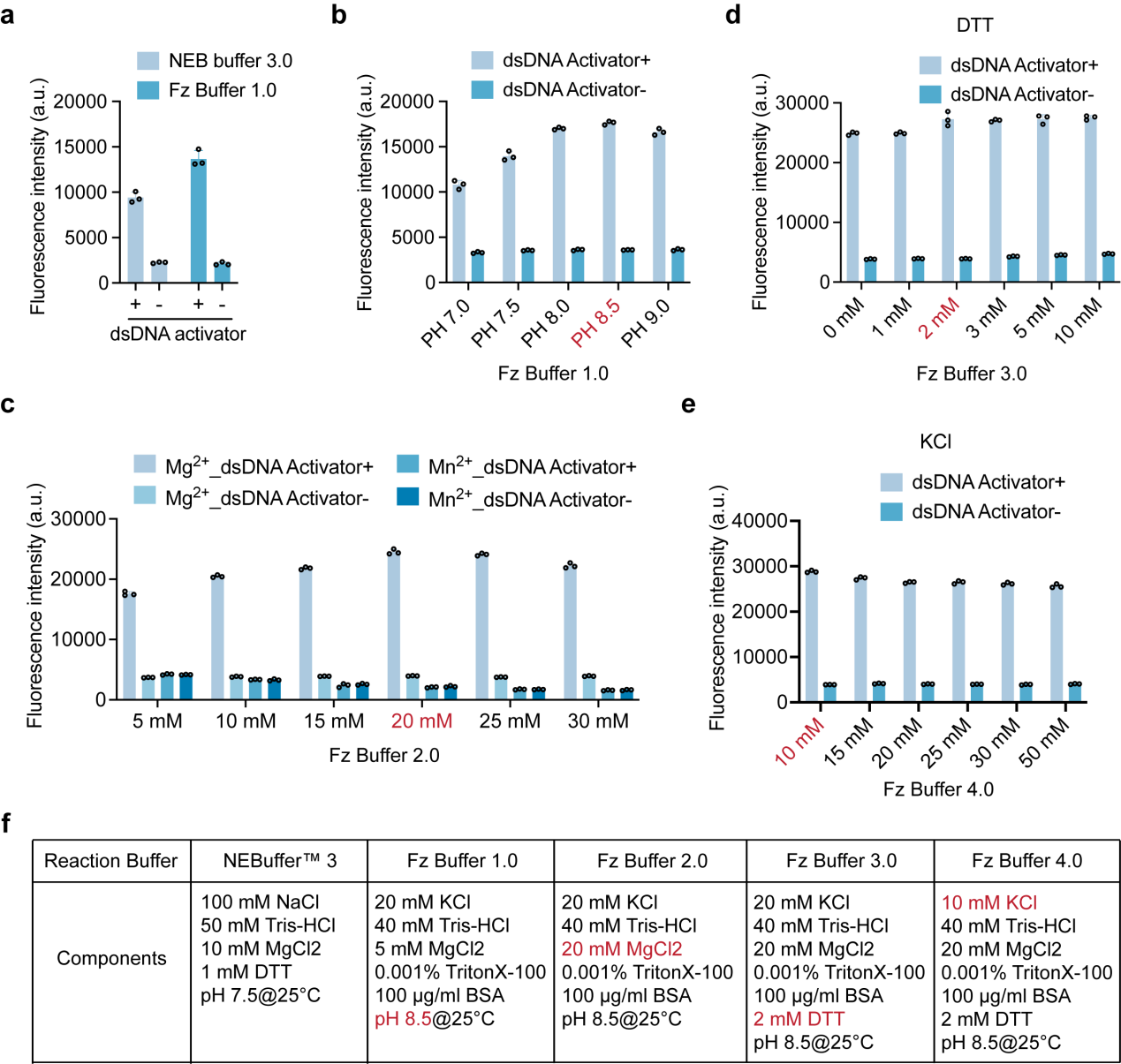

**Extended Data Fig. 2 Optimization of ApmFz2-EP *trans*-cleavage reaction conditions. a**, Comparison of ApmFz2-EP *trans*-cleavage activity in NEBuffer™ 3.0 and Fz Buffer 1.0 using a dsDNA activator. **b–e**, Effects of pH (b), divalent metal ion species and concentration (c), DTT concentration (d), and KCl concentration (e) on ApmFz2-EP *trans*-cleavage activity. **f**, Composition of the buffer systems evaluated during reaction optimization. +, presence of activator template; −, absence of activator template. All data are presented as mean ± s.d. (*n* = 3 technical replicates).

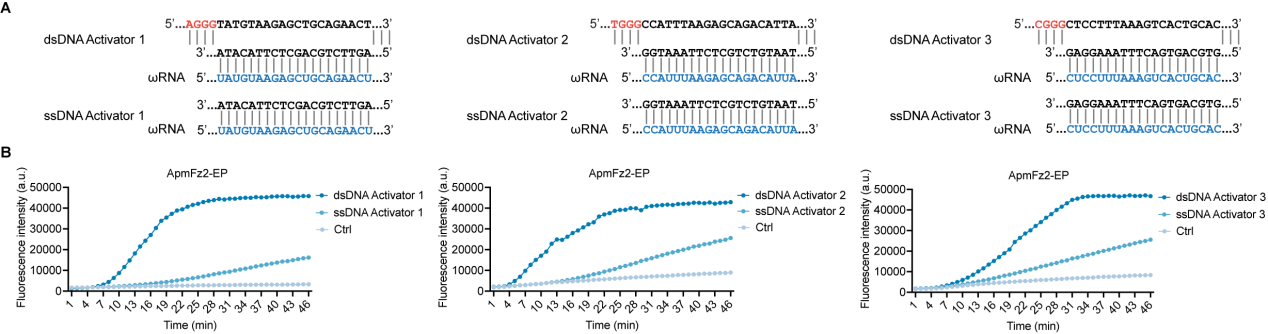

**Extended Data Fig. 3 Comparison of ssDNA- and dsDNA-triggered ApmFz2-EP *trans*-cleavage activity. a**, Sequences of three ssDNA and dsDNA activator pairs and their corresponding guide RNAs. Guide RNA sequences are highlighted in blue, and TAM sequences are indicated in red. **b**, Kinetic analysis of ApmFz2-EP *trans*-cleavage activity triggered by ssDNA and dsDNA activators. Ctrl, no-template control.

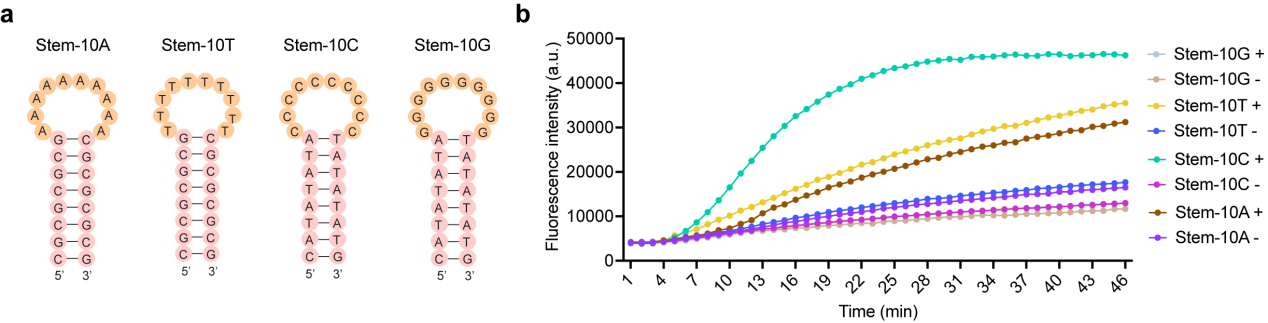

**Extended Data Fig. 4 ApmFz2-EP *trans*-cleavage activity on structured ssDNA reporters. a**, Schematic of ssDNA reporters designed with secondary structures. **b**, Assessment of ApmFz2-EP *trans*-cleavage activity on structured ssDNA reporters in the presence of a dsDNA activator. +, presence of activator template; −, absence of activator template.

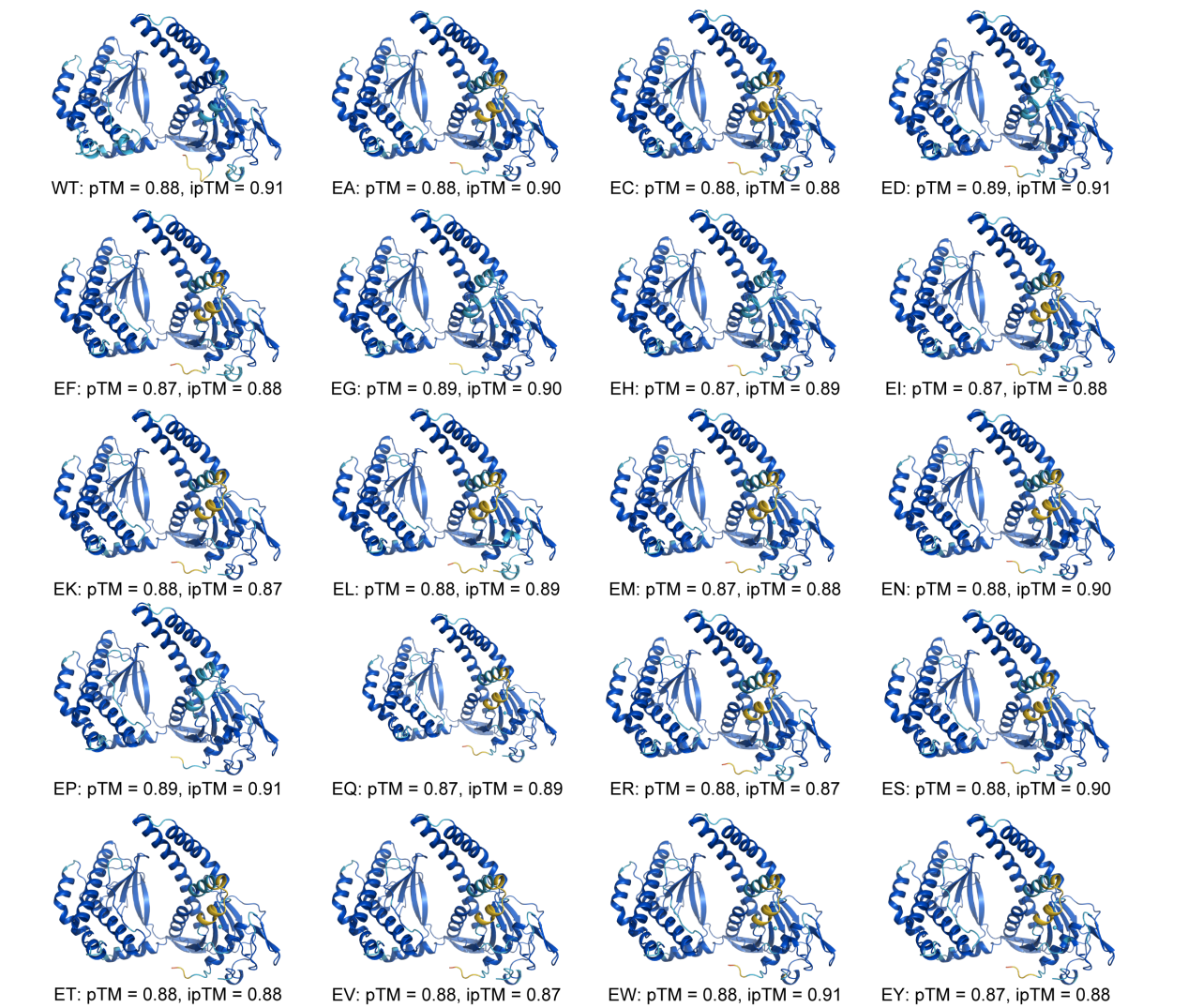

**Extended Data Fig. 5 AlphaFold3 structural models of ApmFz2 double variants.** Predicted structures are shown for wild-type ApmFz2 and experimentally characterized ApmFz2 double variants. Prediction confidence metrics, including pTM and ipTM scores, are indicated for each model. All variants were predicted with high confidence (pTM > 0.85 and ipTM > 0.85). Structures are colored according to the standard AlphaFold confidence scheme, with colors corresponding to residue-level pLDDT values.

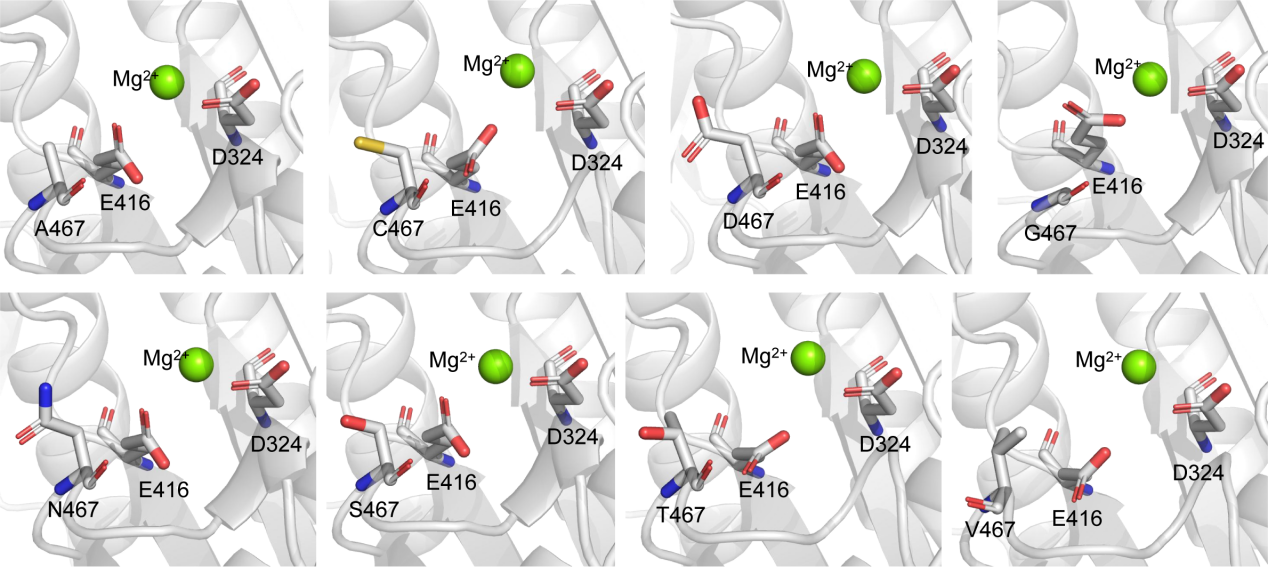

**Extended Data Fig. 6 Structural models of the catalytic centers in ApmFz2 variants with small-to-medium substitutions at position 467.** AlphaFold3-predicted catalytic-center models are shown for ApmFz2 variants containing A467, C467, D467, G467, N467, S467, T467, or V467 substitutions. The engineered E416 residue, catalytic D324 residue, Mg^2+^ ion, and substituted residue at position 467 are indicated.

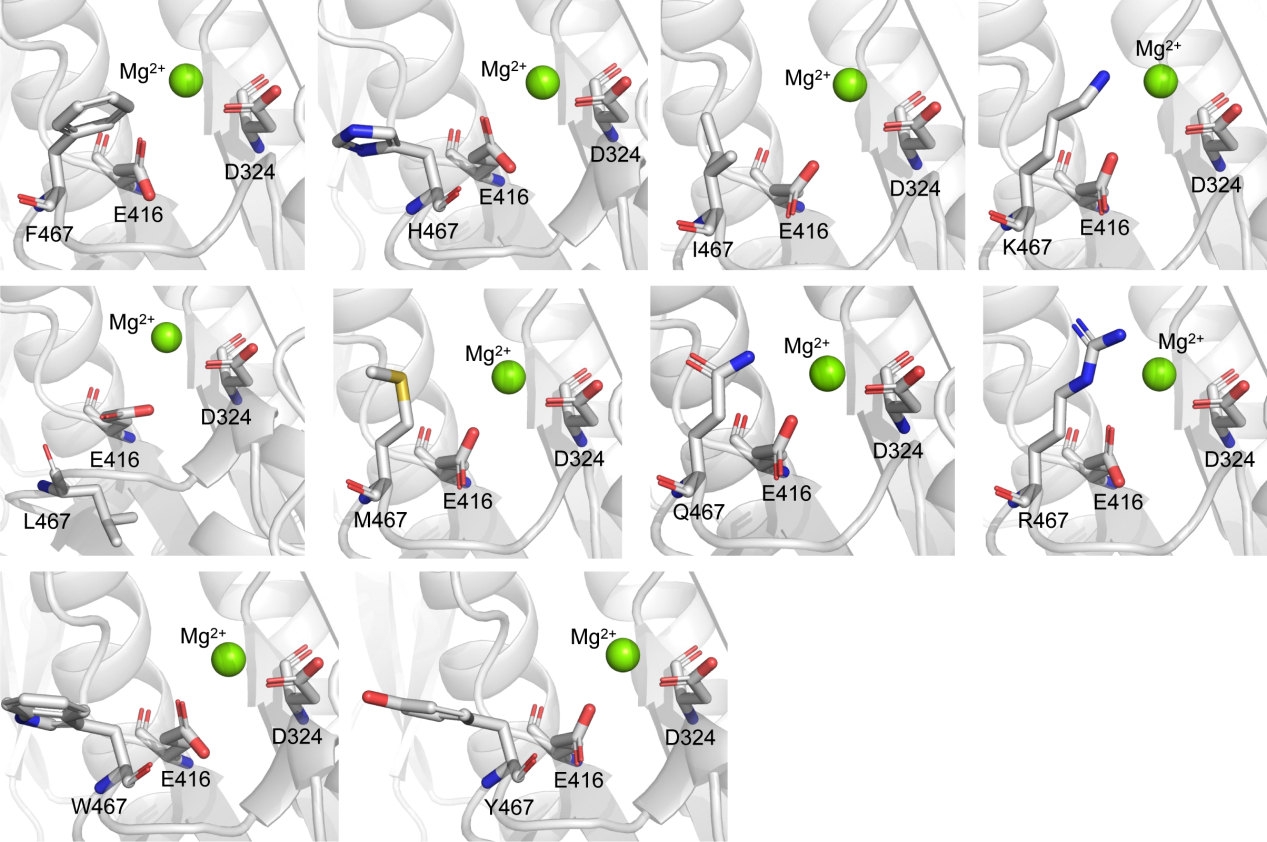

**Extended Data Fig. 7 Structural models of catalytic centers in ApmFz2 variants with bulky substitutions at position 467.** AlphaFold3-predicted catalytic center models are shown for ApmFz2 variants with bulky substitutions at position 467. The engineered E416 residue, catalytic D324 residue, Mg^2+^ ion, and substituted residue position 467 are indicated.

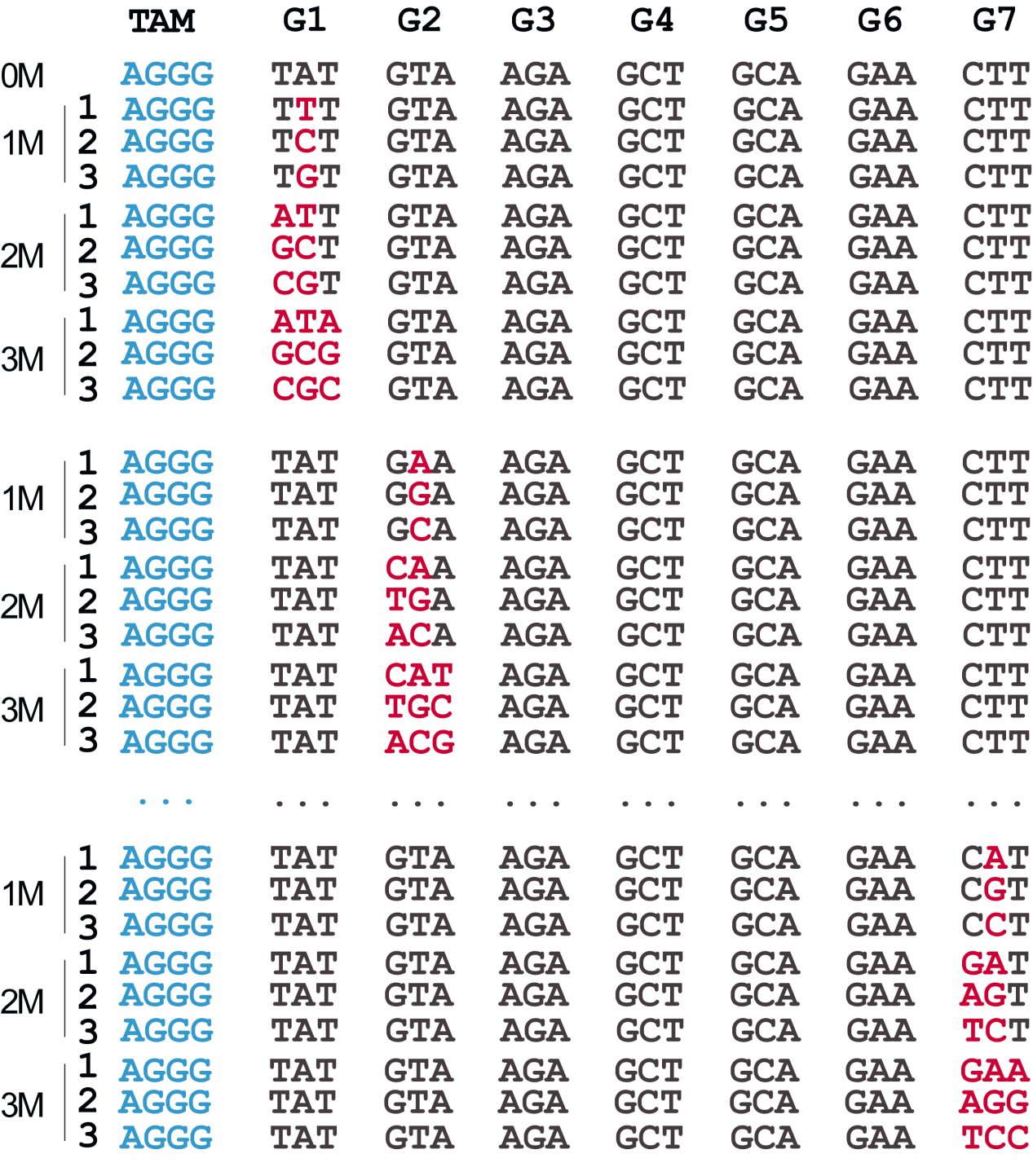

**Extended Data Fig. 8 Schematic of mismatch positions and mismatch numbers within the target sequence.** The 21-bp target region was divided into seven 3-bp groups for mismatch analysis. 0M indicates a fully matched target, whereas 1M, 2M, and 3M indicate targets containing one, two, or three nucleotide mismatches, respectively. Mismatched nucleotides are shown in red.

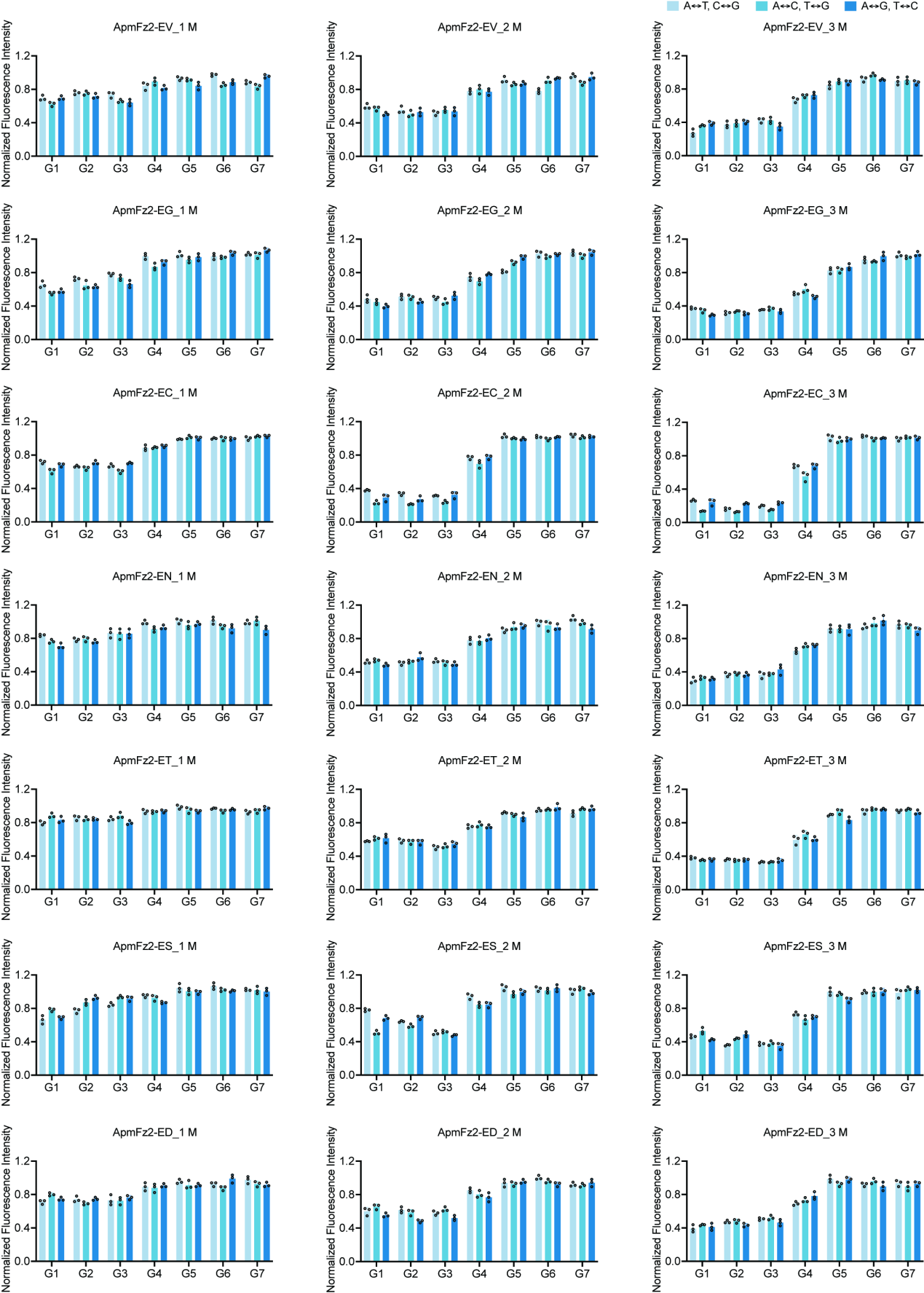

**Extended Data Fig. 9 Mismatch-tolerance profiles of *trans*-cleavage-active ApmFz2 variants.** Normalized fluorescence signals are shown for *trans*-cleavage-active ApmFz2 variants tested against targets containing different mismatch positions and mismatch numbers. 1M, 2M, and 3M indicate targets containing one, two, or three mismatches, respectively. Data are shown as means ± s.d. (*n* = 3 technical replicates).

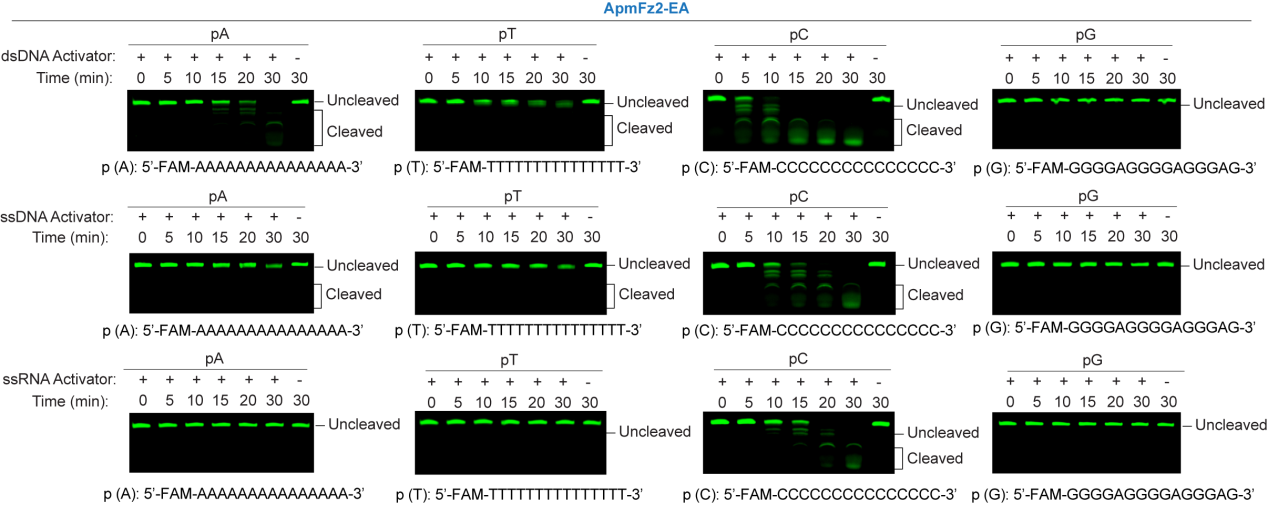

**Extended Data Fig. 10 Reporter sequence preference of ApmFz2-EA *trans*-cleavage activity.** AmpFz2-EA *trans*-cleavage activity assessed using 5′-FAM-labeled poly(A), poly(T), poly(C), and poly(G) ssDNA reporters in the presence of dsDNA, ssDNA, or ssRNA activators, followed by urea-PAGE. p(A), poly(A); p(T), poly(T); p(C), poly(C); p(G), poly(G).

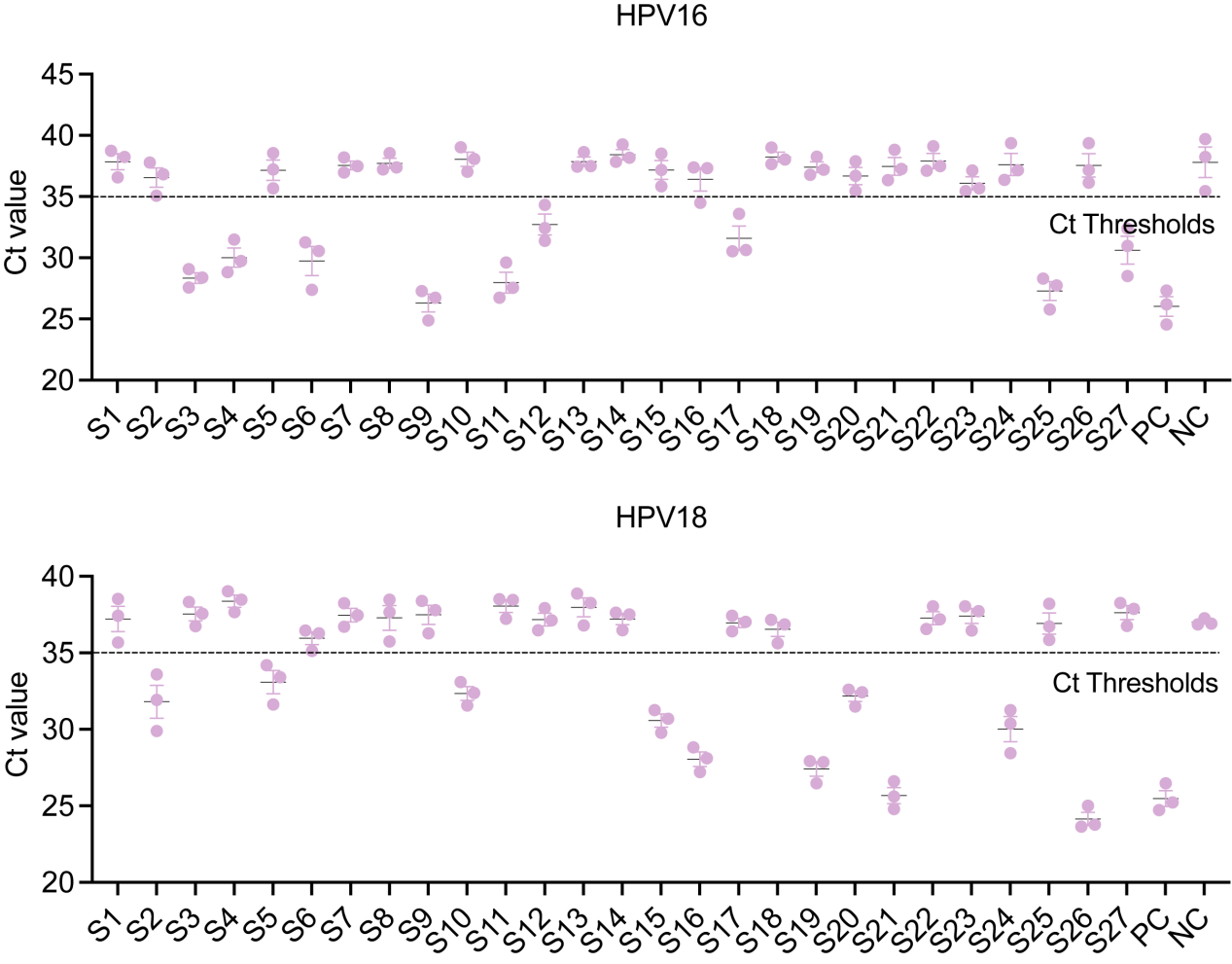

**Extended Data Fig. 11 qPCR validation of HPV16 and HPV18 infection in 27 clinical samples.** HPV16- and HPV18-positive clinical samples were identified by qPCR. PC, positive control containing the pUC57-L1 plasmid template; NC, no-template negative control. Data are shown as means ± s.d. (*n* = 3 technical replicates).

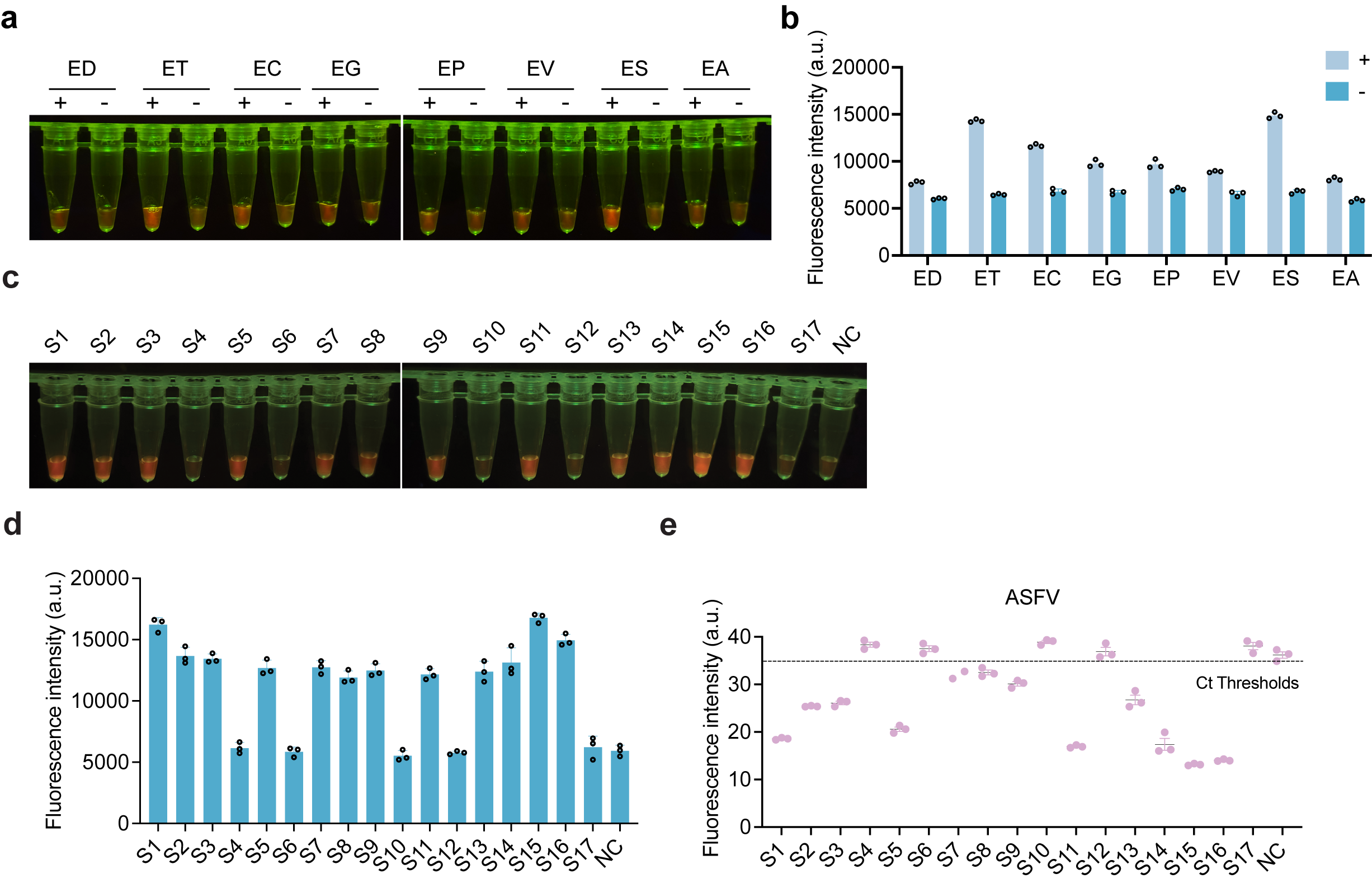

**Extended Data Fig. 12 Evaluation of a one-pot nucleic acid detection using *trans*-cleavage-active ApmFz2 variants. a, b,** One-pot detection assay readouts using blue-light visualization (a) and fluorescence intensity measurements (b). +, presence of pUC57-ASFV-p72 plasmid; −, no-template control. Data are presented as mean ± s.d. (n = 3 technical replicates). **c,d,** Detection of 17 clinical ASFV samples using the FINDER one-pot assay, shown by blue-light visualization (c) and fluorescence intensity measurements (d). Data are shown as mean ± s.d. (*n* = 3 technical replicates). **e**, qPCR analysis of the same 17 clinical ASFV samples used as the reference standard. NC, no-template negative control. Data are shown as mean ± s.d. (*n* = 3 technical replicates).

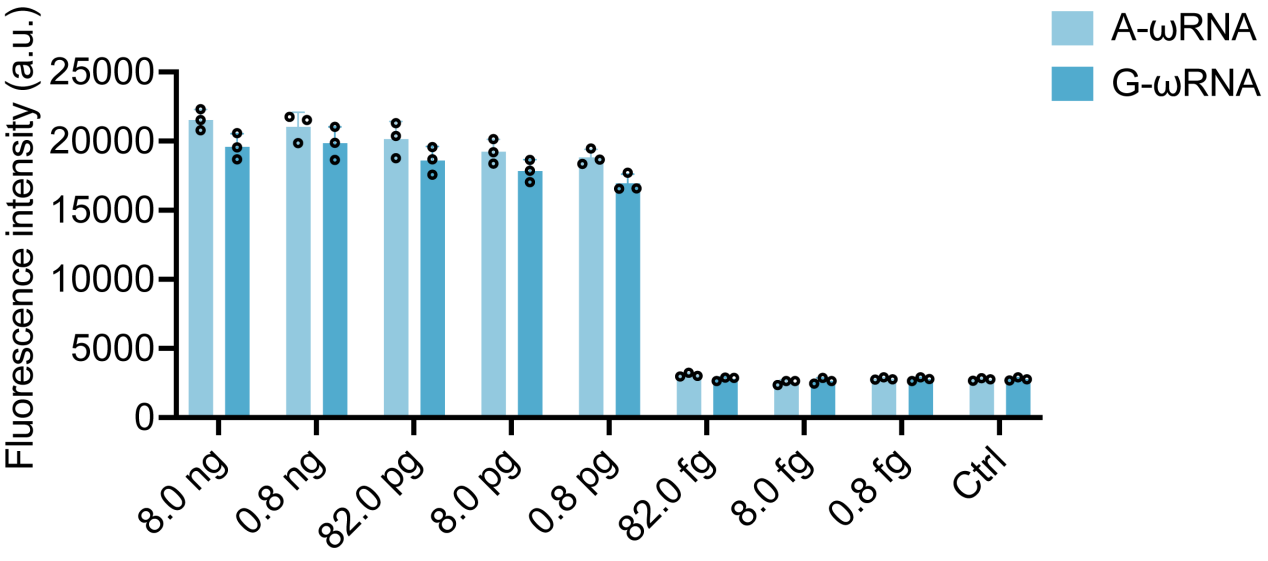

**Extended Data Fig. 13 Sensitivity evaluation of ApmFz2-EA-based SNV genotyping.** Analytical sensitivity of the ApmFz2-EA-mediated FINDER assay for FecB genotyping using sheep genomic DNA as input. Ctrl, no-template control. Data are shown as means ± s.d. (*n* = 3 technical replicates).
